# The distinct role of actin isoforms on mechanosensing-based maturation of hiPSC-derived neurons unveiled by isoform-specific mutations

**DOI:** 10.64898/2026.05.11.724094

**Authors:** Elena Battirossi, Indra Niehaus, Irene Pertici, Giada Magni, Cristiano D’Andrea, Johannes N. Greve, Massimo Reconditi, Nataliya Di Donato, Vincenzo Lombardi, Pasquale Bianco

**Affiliations:** PhysioLab, University of Florence, Florence, Italy; Department of Human Genetics, Hannover Medical School, Hannover, Germany; Cnr-Institute of Applied Physics “Nello Carrara”, Sesto Fiorentino, Italy; Institute for Biophysical Chemistry and Structural Biochemistry, Hannover Medical School, Hannover, Germany

**Keywords:** neuron maturation, neuron mechanosensing, cytoskeletal actin mutations, Dual Laser Optical Tweezers, Baraitser-Winter cerebrofrontofacial syndrome

## Abstract

Neuronal maturation is governed by the integration of intrinsic programs, such as gene regulation and cellular metabolism, with external mechanical cues. The conversion of mechanical forces into developmental responses (mechanotransduction) requires mechanical coupling of neurons to the extracellular matrix and neighboring cells. This coupling is mediated by the actin cytoskeleton and associated transmembrane protein complexes. However, the specific role of the β- and γ-actin isoforms in mechanotransduction, and the functional consequences of their pathogenic mutations, remain poorly understood. To address this question, we combined optical tweezers mechanics with immunofluorescence in human iPSC-derived neural progenitor cells (NPCs) and postmitotic neurons (NCs), both wild-type (WT) and carrying pathogenic mutations in β-actin (R196H) or γ-actin (T203M) associated with Baraitser-Winter cerebrofrontofacial (BWCFF) syndrome. Using membrane tether elongation as a proxy for early protrusion formation, we found that γ-T203M NPCs require a fourfold lower force than WT NPCs and, upon differentiation, fail to develop the neurite-filopodia architecture retaining abundant immature protrusions In contrast, β-R196H NPCs exhibit only reduction of cell surface tension associated with increased neuron fragility, without pronounced protrusion abnormalities. These results reveal non-redundant roles of β- and γ-actin isoforms in neuronal mechanotransduction and provide mechanistic insight into the pathogenesis of BWCFF syndrome.

## Introduction

Maturation of neurons is a complex process that spans from the proliferative stage proper of neural progenitor cells (NPCs) to the postmitotic stage in which neurons (NCs) undergo chemical, structural and functional changes leading to axon guidance, dendrite development and synaptogenesis. The process is controlled by the integration of intrinsic factors, such as gene regulation and metabolism, with extrinsic factors consisting in external stimuli originating from the surrounding environment. These stimuli, which can be chemical, electrical or mechanical, require specific transducers on the cell membrane and acquire progressively more importance throughout maturation.

Neuron mechanosensing is the ability of the neuron to detect mechanical stimuli and transduce them into developmental, biochemical and structural responses. Mechanical stimuli may originate either from passive forces applied by the surrounding tissue acting as a plastic scaffold or from the active forces exerted by the cell itself during specific interactions with the extracellular matrix (ECM) or a target cell and involve an efficient mechanical coupling between the intracellular actin cytoskeleton and the extracellular partner(s).

During neuronal development, these forces change their nature depending on the stage of maturation (for an extensive review, see^1^). The immature neurite, the axonal process constituted only by microfilaments (actin cytoskeleton)^2^, advances thanks to the combination of the force arising from actin polymerization at the leading edge and the force resulting from actin-myosin interaction, both supporting the retrograde actin flow. Given the binding of actin cytoskeleton to the substratum via the cell adhesion molecular complex, the retrograde actin flow exerts a traction force on the substratum, resulting in elongation of the leading edge. The connection of the neurite with the target cell induces a new tensile force that is maintained by the increasing distance between the two cells (stretch growth^3,4^). Tension within the axon promotes its maturation, with the appearance of mitochondria and microtubules, and inhibits the growth of collateral neurites^5,6^. By reproducing this stretch growth *in vitro* with the remotely controlled manipulation of interiorized magnetic nanoparticles^6^, it has been directly demonstrated that the chronic maintenance of a neurite under tension accelerates its elongation and maturation.

Forces generated by cell environment and cell-cell interactions are responsible for the mechano-transduction cascade of translational, chemical and structural steps leading to neuronal maturation, depending on the efficient mechanical coupling of the soma throughout the neurite to the substratum or the target cell. The structural elements underpinning this coupling are the actin cytoskeleton and the protein complex that connects it to the ECM or the target cell.

The use of disease-specific mutants of either primary embryonic neurons or neurons derived from human induced pluripotent stem cells (iPSCs) that mimic the maturing tissue, combined with immunofluorescence imaging, has opened new possibilities for understanding both the role of each molecular player in the coupling system and the loss-of-function associated with its causative mutation. In cultured mouse hippocampal neurons, this approach allowed the detailed description of the molecular basis of the conversion of the retrograde flow of polymerizing actin into leading-edge advancement and, through selective knock-out, the identification of the protein that bridges the actin cytoskeleton to the transmembrane protein^7^.

A similar integrated approach of mechanics and immunofluorescence imaging is used here to establish the role of cytoskeletal actin isoforms in the mechanosensing-based maturation of human iPSC-derived neurons. Two actin isoforms are present in the cytoplasm of neurons, β-actin and γ-actin (encoded by the *ACTB* and *ACTG1* gene, respectively). β- and γ-actin differ by only four amino acids but exhibit distinct polymerization kinetics, filament stability and interactions with actin-binding proteins (ABPs) in the submembrane cytoskeleton, the cortex controlling the balance between cell surface protrusion and stabilization^8–16^. However, the role of β-actin and γ-actin in NPCs differentiation is not known.

Mutations in the *ACTB* and *ACTG1* genes cause non-muscle actinopathies (NMAs), a heterogeneous group of disorders including the neurodevelopmental Baraitser-Winter cerebrofrontofacial syndrome (BWCFF), which is the most prevalent and is characterized by cortical malformations, intellectual disability and craniofacial anomalies ^17,18^. β-R196H and γ-T203M are two of the most frequent actin variants leading to BWCFF. Both disrupted residues are essential for actin polymerization, actin-ABP interactions, filament and cortex stability and neurite extension^19–22^. Both substitutions affect residues located in subdomain 4 of the actin molecule, which is crucial for stabilizing intersubunit contacts within F-actin and for mediating interactions with ABPs^23^. Since patients carrying the same aminoacidic substitution in both isoforms have not been observed, these variants represent the closest clinically relevant models for investigating whether alterations of neuron mechanics caused by mutations in this region depending on the isoform. Because mutations in either isoform are heterozygous missense substitutions and the mutant protein remains stable, the resulting cellular phenotype extends beyond a simple loss-of-function mechanism^20,21^. Therefore, this system can be used to define isoform-specific blunting of mechanotransduction during neuronal maturation and to provide mechanistic insight into the neurodevelopmental etiology of the disorder.

To this scope, we combine immunofluorescence with dual-laser optical tweezers (DLOT) technology^24,25^ to characterize the morphological and mechanical properties of human iPSC-derived WT and β- and γ-actin mutant neuronal cells at developmental stages of NPCs, at 18-25 days after iPSC seeding and NCs, at day 14 after transfer to maturation medium. The DLOT allows quantitative estimates of the mechanical parameters of interest. The protocol provides that the cell is brought into contact with a bead trapped in the focus of the DLOT and pulled to elicit the formation/elongation of a tether, a 100-200 nm diameter cylindrical protrusion that is progressively invaded by the actin cytoskeleton^26^. This process is particularly rapid in neuronal cells^27^. It is characterized by a transition in tether mechanical properties, from an initially viscoelastic response, arising from viscous drag between the two membrane leaflets and, especially, between the inner leaflet and the cytoskeleton^28^ modulated by membrane-cortex adhesion, to a dynamically stable, actin-containing protrusion of the cell surface. This structure resembles those naturally occurring with the development of filopodia^29^ and tunnelling nanotubes^30,31^. Thus, tether dynamics provides a controlled assay of cell-surface mechanics and of mechanosensing-dependent developmental plasticity.

The functional phenotype of WT, β- and γ-actin mutant neurons at the two developmental stages is identified by viscoelastic parameters that characterize cell-surface resistance to tether formation and elongation, including the effective stiffness responsible for the quasi-static tension that assists actin recruitment within the tether. Compared with WT cells, γ-T203M NPCs show a four-fold reduction in effective stiffness versus tether elongation, similar to the drop induced by the actin polymerization inhibitor Latrunculin-A. Imaging analysis reveals that the soma of the γ-mutant NPCs misses the smooth contour typical of the WT NPCs under the tensing effect of interneuron connections and appears fringed and sprinkled with randomly curved protrusions. During maturation, γ-mutant NCs maintain numerous undifferentiated protrusions, failing to evolve towards a reduced number of normal neurites with filopodia characteristic of the WT NCs architecture. Neither of these mechanical and morphological differences is observed in the β-actin R196H mutant, which instead exhibits reduced cell surface tension, associated with marked neuron fragility.

## Results

### WT and β- or γ-actin mutant cells display distinct morphologies and sizes at both NPC and NC stages

The cell differentiation timeline is summarized in the upper panel of Fig. 1. Human iPSCs (day 0) progress through embryoid body (day 5) and neural rosette organization (days 11–12), leading to the generation of neural progenitor cells (NPCs, day 18). Upon neuronal induction (from days 25-28) and maturation (from days 33-36), cells progressively acquire elongated processes and increased neurite complexity, culminating in mature neuron-like cells (NCs, days 47–50, 14 days after seeding in maturation medium). Mechanical and immunofluorescence characterization are performed at the NPC stage (day 18) and at the NC stage (days 47–50), as indicated by the arrows.

**Fig. 1.**
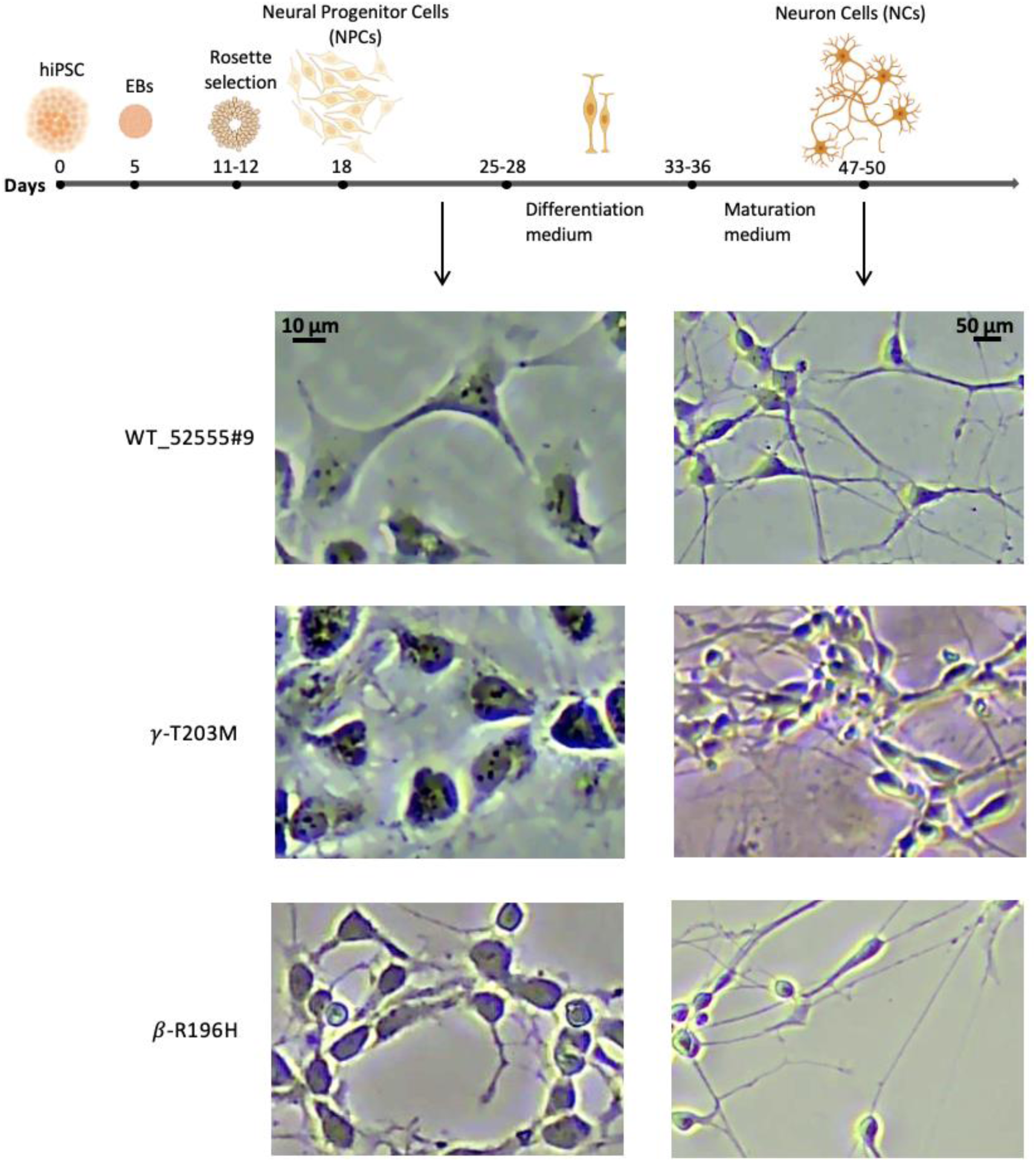
Differentiation protocol and morphometric characteristics of human iPSC-derived neurons at NPC and NC stages. **Upper panel**. Time course (days) of *in vitro* differentiation from human iPSC through embryoid bodies (EBs), rosette selection, NPC expansion (from day 18) and terminal NC differentiation (from days 25-28), and maturation (from days 33-36). **Lower panel**. Representative bright-field images of NPCs (left column, between days 18 and 28) and NCs (right column, after 14 days in the maturation medium). Upper row, WT line 52555#9; middle row, γ-T203M mutant; lower row, β-R196H mutant. Scale bars: 10 µm (NPCs) and 50 µm (NCs).

Bright-field microscopy analysis preliminary to mechanical investigation (Fig. 1, lower panel) shows that, in agreement with the differentiation-dependent increase in neuronal size^32^, WT NCs have larger somas than WT NPCs. A semi-quantitative analysis (see Materials and Methods and^33^) confirms a significant increase in size with differentiation (Table 1): the area of WT NPCs is ∼ 160 µm^2^ and is 4 times larger in WT NCs (∼ 650 µm²). WT values were calculated as the mean of all WT data points pooled together, without distinguishing the line of origin (see Materials and Methods and Table S1). Among mutant cells, while the γ-T203M mutant displays a soma size similar to WT cells at either stage of differentiation (p > 0.4), the β-R196H mutant at the NPC stage shows a reduced soma area (∼ 100 µm^2^, −40%, p < 0.01). Upon maturation, the β-R196H NCs also increase their size, attaining a value not significantly different from the other NCs (p ≥ 0.4).

**Table 1.** Morphometric analysis of neuron soma at the two stages of differentiation selected for the study.

| Experimental model | | *N* | *n* | Area ($\mu\text{m}^2$) |
| --- | --- | --- | --- | --- |
| NPC | WT | 6 | 528 | 165 ± 22 |
| NPC | γ-T203M | 3 | 246 | 164 ± 44 |
| NPC | β-R196H | 4 | 212 | 103 ± 14 <sup>#</sup> |
| NC | WT | 10 | 425 | 648 ± 96 |
| NC | γ-T203M | 5 | 235 | 675 ± 130 |
| NC | β-R196H | 5 | 245 | 650 ± 150 |

Soma size was quantified from bright-field images using Fiji/ImageJ software. *N* is the number of images used for the analysis; *n* is the number of cells analyzed. <sup>#</sup> indicates a significant difference (*P* < 0.05) with respect to wild type.

### Mechanical characteristics of tether formation and elongation in WT NPCs

The protocol described here for the mechanical analysis of the WT NPCs from 52555#9 line is the prototype for the analysis conducted across all experimental models. The cell is brought into contact with the trapped bead (Fig. 2A, phase 0, initial position) and then preliminarily pushed against the bead (phase 1) to establish a stable bead-membrane bond. Tether extraction/elongation is obtained by moving the cell away with a ramp-faced pull (phase 2, velocity 1-8 μm s^−1^) of amplitude ranging from 10 to 50 μm. At the end of the ramp the position is held for 10-30 s (phase 3), and then the cell is returned to the initial position (phase 4). After a 15-100 s pause (phase 5), the cycle is repeated (phases 6–8). The protocol was generally repeated over multiple cycles to further probe the mechanical response. The duration of the pause between cycles was randomly assigned at each cycle, with long pauses (∼100 s) used less frequently, as longer waiting times tend to reduce measurement stability due to thermal fluctuations of the DLOT.

**Fig. 2.**
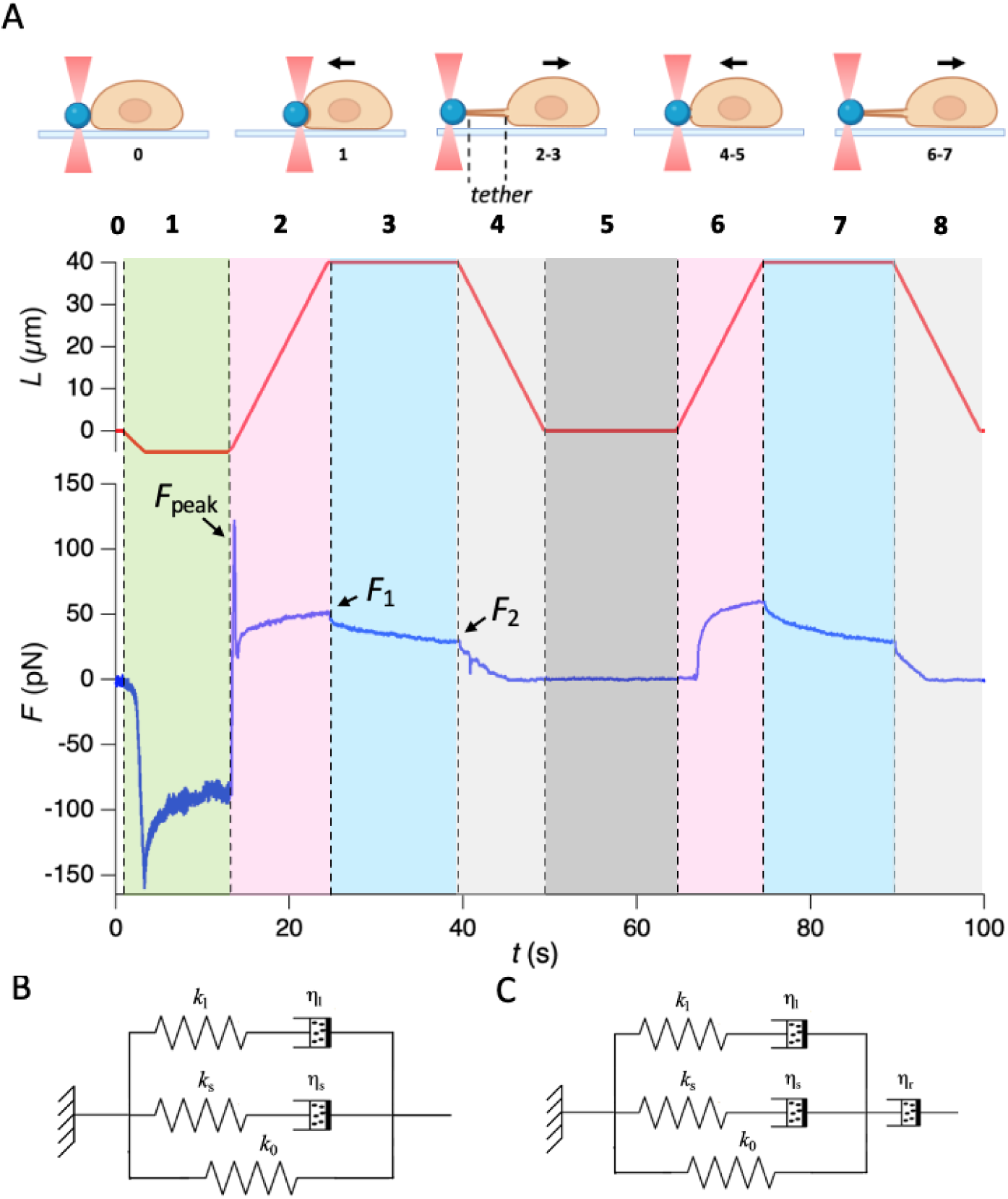
Mechanical characteristics of tether extraction and elongation. **A. Upper panel.** Schematics of the mechanical protocol applied by controlling the position of the stage carrying the cell. Displacements indicated by arrows are imposed parallel to the plane on which the cell lies. Numbers indicate the corresponding phases of the mechanical maneuvers as shown in the lower panel. **Lower panel.** Force response (*F*, blue trace, positive in response to elongation) to the imposed displacement (*L*, red trace). Vertical dashed lines mark transitions between phases marked by specific colors and numbers. In the first cycle, the cell is brought into contact with the bead (initial position, phase 0) and then pushed against it to establish a stable bead-membrane interaction (phase 1). Tether extraction/elongation is obtained by moving away the cell with a ramp-shaped elongation at velocity 4 μm s^−1^ (phase 2) and amplitude 45 μm. At the end of the ramp the position is held for 15 s (phase 3) and then the cell is returned to the initial position (phase 4). After a 15 s pause (phase 5) the cycle is repeated (phases 6–8). Background colors to identify the phases numbered as in the upper panel: (0) white, (1) light green, (2,6) pink, (3,7) light blue, (4, 8) light gray, (5) gray. The force trace reveals three characteristic signatures: *F*_peak_, the early force peak at the onset of first elongation; *F*_1_, the force attained at the end of the elongation; *F*_2_, the quasi-steady force attained upon relaxation at the end of the hold. **B.** Equivalent mechanical model (Zener model) describing the viscoelastic nature of the responses to the lengthening-shortening cycle. The model comprises three elastic elements (*k*₀, *k*_s_, *k*_l_) and two dashpots (η_s_, η_l_) as detailed in the text. **C.** Mechanical model extended to include an additional dashpot (ηᵣ) in series to the Zener model, to account for the relaxation that, according to the literature, occurs in the time scale of minutes (see text).

In the first cycle, the force response to the pull is characterized by an early rapid rise to a peak (*F*_peak_) followed by an abrupt drop, and a subsequent rise at a rate that progressively decreases up to a constant value that, at the end of the ramp, brings the force to a maximum (*F*_1_). During the hold, the force relaxes to a quasi-steady value *F*_2._ Between the first and the second cycle the pre-existing base for tether formation is preserved, allowing elongation without the *F*_peak_ response. This clearly indicates that the *F*_peak_ response marks the initial abrupt membrane-cortex detachment accompanying tether formation in the relatively high-velocity regime chosen to maximize the resolution of the dynamic and quasi-static force responses (*F*_1_ and *F*_2_, respectively) to tether elongation. In fact, these velocities greatly exceed the physiological membrane extension or filopodial/axonal growth rates (< 0.05 µm s^−1^)^34^ but they allow the system to rapidly enter a mechanical regime suitable for quantitative optical-trapping measurements of cell-surface dynamics. The following analysis focuses on the response characteristics of the subsequent cycles of perturbations, thus excluding the early, unphysiological *F*_peak_ response.

The force relaxation recorded during the hold is a manifestation of the passive viscoelastic nature of the cell surface’s resistance to tether elongation^26–28,35,36^. The relaxation time course is biphasic, according to the equation:

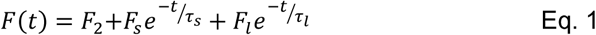

where *F*_s_ and *F*_l_ represent the amplitudes of the short and long relaxation processes, defined by the corresponding time constants, τ_s_ (200-300 ms) and τ_l_ (∼5 s, thus the long process is almost complete within the hold time), and *F*_2_ is the amplitude of the asymptotic force, estimating the quasi-steady force attained at the end of the hold (Table S2). The biexponential relaxation supports an equivalent mechanical model like a Zener model (Fig. 2B), consisting of an elastic element (*k*₀), responsible for the steady force *F*_2_, in parallel with two Maxwell elements, each composed of an elastic spring (*k*_s_ and *k*ₗ) in series with a dashpot (η_s_ and ηₗ, respectively), responsible for the drag resistance opposed to elongation by the elements of the cell surface undergoing reciprocal slippage during the pull ^28^, among which the most relevant is the membrane-cortex system.

The overall amplitude of the relaxation within the 10-30 s of the hold (*F*_s_ + *F*ₗ) is a fraction of the force recovered during the lengthening. According to the literature, further recovery occurs on a much longer time scale (minutes), consistent with the time taken by recruitment of cytoskeletal actin within the tether and its consequent remodeling into a stable protrusion ^26,27,36^. In fact, even if we have not followed the time course of actin recruitment, we find that in all our experimental models (WT, β-R196H and γ-T203M mutant NPCs and NCs) actin entered the tether within 5 min at most after tether extraction (see Materials and Methods and Fig. S1). The small delay taken by the force to rise with elongation in the second cycle (phase 6 in Fig. 2A) indicates that the stage movement to recover the initial position (phase 4) represents an excess with respect to the shortening required to recover the baseline force, suggesting that tether remodeling with actin recruitment has started within the 20 s of the preceding hold with high tether tension (phase 3). The suppression of the tension during the subsequent pause (phase 5) reverses the remodeling process (see^27^ for an extensive review). In fact, as shown in Fig. S2, the delay at the start of elongation in the subsequent cycle is suppressed if the pause is raised from ∼20 s (between 1^st^ and 2^nd^ cycle) to ∼100 s (between 2^nd^ and 3^rd^ cycle). Notably, it is the quasi-static force *F*_2_ which, by itself, triggers the actin recruitment acting as a *vis a fronte*, that induces further force recovery in the time scale of minutes. This process can be simulated with an equivalent mechanical model of the cell surface by introducing a dashpot with a much higher drag coefficient in series with the Zener model (η_r_, Fig. 2C, see also^35^). The analysis hereafter preserves the almost pure viscoelastic response to elongation (depicted by the model in Fig. 2B) by setting the pause preceding every elongation cycle to ∼100 s.

For any given lengthening velocity, both *F*_1_ and *F*_2_ increase with the extent of elongation (Fig. 3A), as expected from the contribution of the elastic elements of the equivalent mechanical model. Instead, for any given elongation, *F*_1_ increases with the elongation velocity while *F*_2_ remains constant (Fig. 3B), as expected from the contribution of the viscous elements, provided that, at the end of the hold, the viscous elements have completely relaxed. This is true assuming that, within the ∼20 s of the hold, the recruitment of actin within the tether is small enough for its contribution to *F*_2_ to be ignored^26,27,36^.

**Fig. 3.**
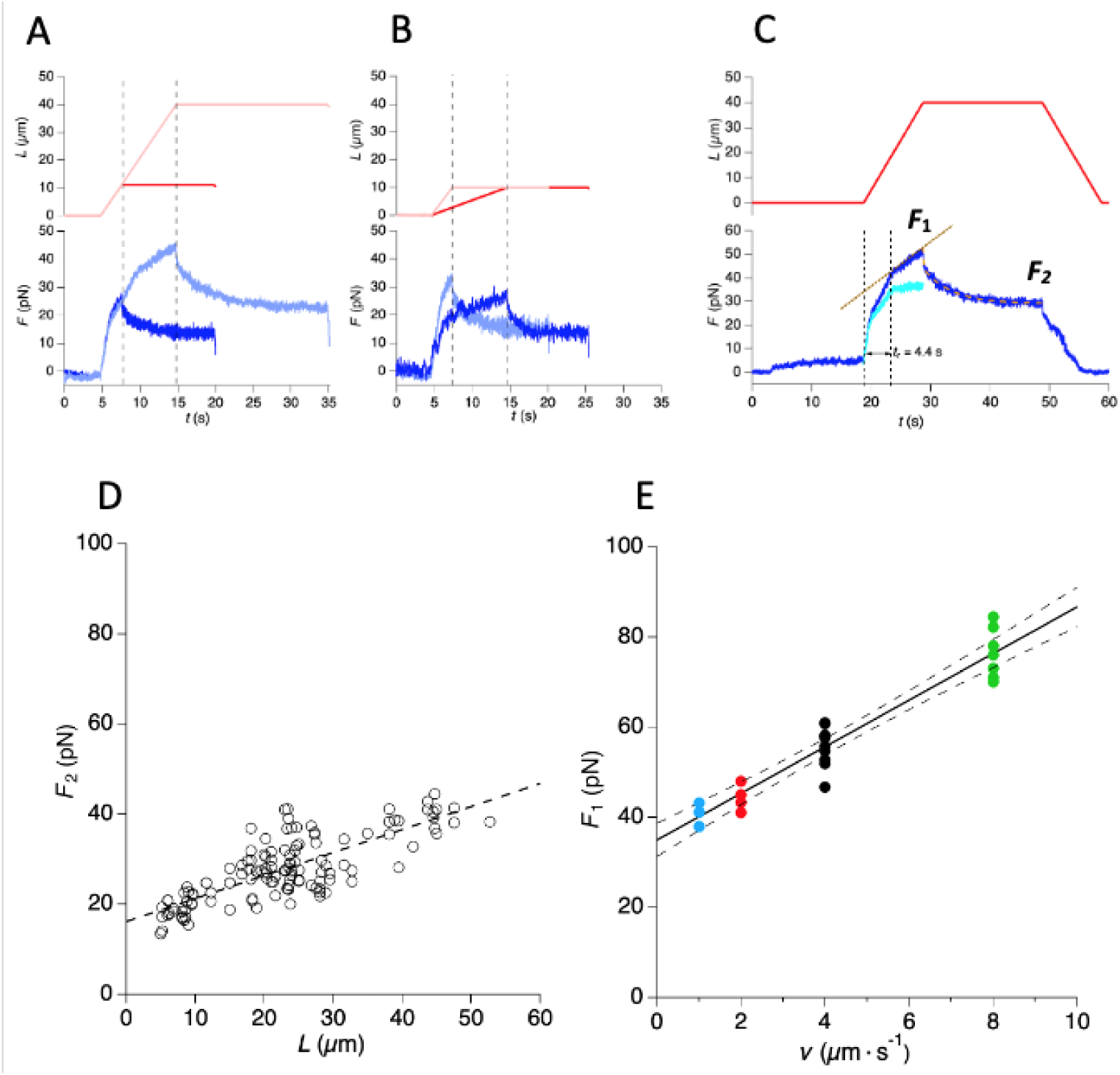
Force response during tether elongation and its dependence on elongation amplitude and velocity. **A.** Superimposed force responses (*F*, blue traces) to two elongations of different amplitude (*L*, red) at velocity 4 µm s⁻¹. *L* = 10 µm, bright red, and 40 µm, light red; bright blue and light blue, *F* for short and long *L*, respectively. Vertical dashed lines mark the time of ramp-hold transition. **B.** Superimposed force responses to two elongations of 10 µm at different velocity (*v*). *v* = 1 µm s⁻¹, bright red, and 4 µm s⁻¹, light red; bright blue and light blue, *F* for smaller and larger *v,* respectively. **C.** Force response (blue) to elongation-hold-shortening cycle (red) of 40 μm at *v* = 4 µm s⁻¹. *F*_1_ and *F*_2_ mark the force at the end of the elongation ramp and at the end of the hold, respectively. Force relaxation during the hold is fit with Eq. 1 (orange dashed line). By subtracting the line tangent to the final part of force response (brown dashed line), the viscoelastic component of the force response to lengthening can be isolated (cyan trace), from which the rise time *t*_r_ (the time required for the force to rise from 10% to 90% of its maximum value, vertical dotted lines) is determined. **D.** Dependence on tether elongation (*L*) of *F*_2_. The dashed line is the linear fit to *F*_2_-*L* relation. **E.** Dependence on *v* of *F*_1_ points collected for elongations lasting longer than *t*_r_ (see Fig. S3). *v* is identified by the color code as in Fig. S3: 1 µm s^−1^ (light blue), 2 µm s^−1^ (red), 4 µm s^−1^ (black) and 8 µm s^−1^ (green). The slope of the relation, estimated by the fit to data with Eq. 3 (continuous line), corresponds to 2πη_eff_ and provides a value for the effective viscosity η_eff_ of 0.81 ± 0.1 pN · s μm^−1^. The ordinate intercept (34.9 ± 3.7 pN) provides an estimate of *F*_2_ for *L* = 40 μm and is in quite good agreement with the value from the *F*_2_-*L* relation in **D** (37.4 ± 3.7 pN for *L* = 40 μm).

The viscoelastic contribution during elongation was isolated by subtracting the final linear elastic component (Fig. 3C, brown dashed line) from the force response (blue trace).The viscoelastic response isolated in this way (cyan trace) reflects the contribution of the Maxwell elements in parallel in the equivalent mechanical model (Fig. 2B). Its rise time, *t*ᵣ, defined as the time from 10 to 90% of the force plateau value, 4.4 s in the WT NPC analyzed here, indicates the minimum elongation time required for the force response to reach the level set by viscous resistance. For any given elongation velocity *v* (1-8 µm s^−1^), once the elongation durations exceeded *t*ᵣ, *F*_1_ increased linearly with tether length (*L*, Fig. S3, filled circles), with a slope that estimates *k*₀. *k*₀ can be estimated also by the slope of the relation between *F*_2_ and *L* (open circles in Fig. S3 and Fig. 3D). The first order regression equation (dashed line) fitted to the *F*_2_ – *L* relation

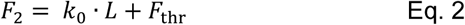

gives a *k*_0_ of 0.53 ± 0.03 pN μm^−1^ (Table S3) and an ordinate intercept, which represents the threshold force *F*_thr_ necessary to induce tether formation of 15.7 ± 0.9 pN, Table S3).

The *F*_1_-*L* relation at an elongation time > *t*_r_ is progressively shifted upward by the increase in *v* from 1 to 8 μm s^−1^ (Fig. S3), underpinning a linear relation between *F*_1_ and *v* (Fig. 3E, where the different colours mark the different velocities: cyan 1, red 2, black 4 and green 8 μm s^−1^, for the details see Fig. S3 legend). The fit of the relation with the first order regression equation),

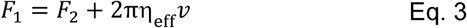

defines the effecti*v*e viscous coefficient for tether elongation, ηₑ_ff_ = 0.81 ± 0.13 pN s μm⁻¹ (Table S3).

### Comparative analysis of the mechanical parameters describing tether dynamics in the different experimental models

To improve the reliability of the comparison with data collected from the two patients with Baraitser-Winter cerebrofrontofacial (BWCFF) syndrome, we integrated the analysis concerning WT NPCs from 52555#9 line with the data extracted from a second WT (SCTi003-A; see Materials and Methods). As detailed in Tables S1 and S3, all the parameters considered in the analysis excluded any significant difference between the two lines. Therefore, hereafter, data from the two WT lines are pooled and averaged for comparison with mutant cells, irrespective of line origin.

Averaged *F*_2_-*L* data of NPCs from the two WT lines (Fig. 4A, black circles) are superimposed on the mean data obtained from β-R196H NPCs (blue circles), γ-T203M NPCs (green circle) and Latrunculin-A treated WT NPCs (magenta circles). Lines show the fit of Eq. 2 to the data, identified by the same color code. The slope of the relation (the quasi-static stiffness *k*_0_) of WT NPCs is quite similar to that of β-mutant NPCs, while it is significantly lower in both γ-mutant NPCs and Latrunculin-A-treated NPCs. The ordinate intercept of the relation (the threshold force for tether formation *F*_thr_), instead, appears reduced in β-mutant NPCs but not in γ-mutant NPCs, and is even more dramatically reduced in Latrunculin-A treated NPCs. The parameters estimated from the linear fit of *F*_2_-*L* data across all experimental models at the NPC stage, along with the statistical significance of their differences, are reported in Tables 2 (*F*_thr_) and 3 (*k*_0_). *k*_0_ (0.52 pN µm^−1^ in WT cells, Table 2), is not significantly affected in the β-R196H mutant, while it is reduced in γ-T203M mutants, to a value (0.12 pN nm^−1^) not significantly different to that of Latrunculin-A treated NPCs. As far as *F*_thr_ (Table 2), it does not show a significant reduction in the γ-T203M mutants, while it drops to ∼0.6 the WT value (p < 0.0001) in the β-R196H mutant. An even larger reduction of *F*_thr_ (to ∼0.1 the WT value) is induced in Latrunculin-A-treated NPCs.

**Fig. 4.**
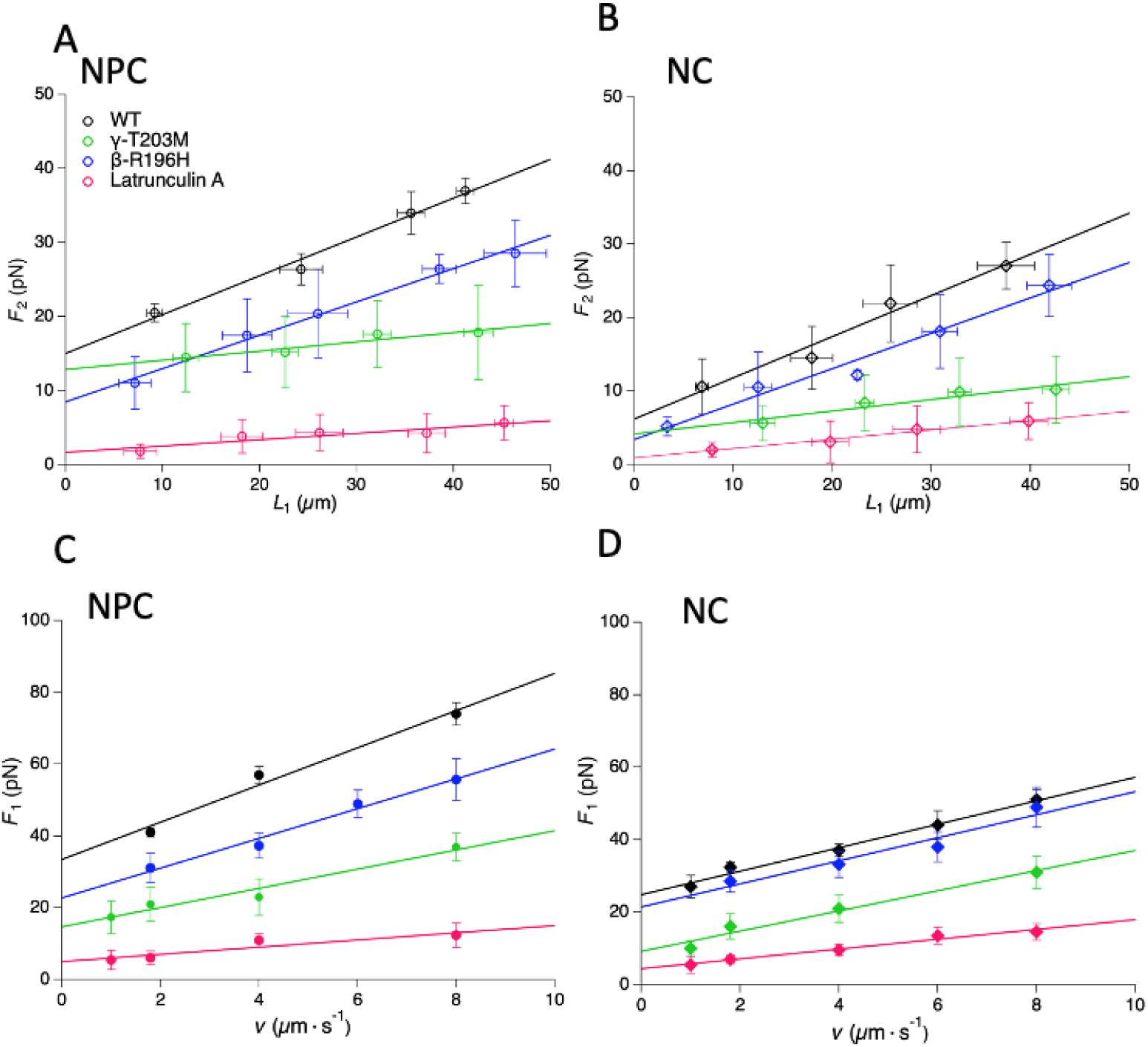
Comparative analysis of *F*_2_-*L* and *F*_1_-*v* relations among all experimental models. **Upper panels.** *F*_2_-*L* relations at NPC (**A**) and NC (**B**) stages. The color of symbols (means ± SD) and lines (fit) identifies the experimental model: black, WT; blue, β-R196H mutant; green, γ-T203M mutant; magenta, Latrunculin-A-treated cells. **Lower panels.** *F*_1_–*v* relation at NPC (**C**) and NC (**D**) stages. Color of symbols (means ± SD) and lines (fit) as in the upper panels. The parameters extracted from the fits are reported in Tables 2 and 3.

**Table 2.** Mean ± SD of tether extraction and elongation parameters for all experimental models.

| Experimental models | | $n$ | $F_{\text{thr}}$ (pN) | $k_0$ (pN/$\mu\text{m}$) | $n_1$ | $\eta_{\text{eff}}$ (pN·s/$\mu\text{m}$) |
| --- | --- | --- | --- | --- | --- | --- |
| NPC | WT | 129 | $14.9 \pm 0.8$ | $0.52 \pm 0.03$ | 34 | $0.83 \pm 0.11$ |
| NPC | γ-T203M | 73 | $12.9 \pm 0.9$ <sup>\$</sup> | $0.12 \pm 0.03$ <sup>#</sup> | 20 | $0.45 \pm 0.14$ <sup>#</sup> |
| NPC | β-R196H | 91 | $8.5 \pm 0.8$ <sup>\$, #</sup> | $0.45 \pm 0.03$ <sup>\$</sup> | 47 | $0.65 \pm 0.19$ <sup>\$</sup> |
| NPC | 52555#9 + Latr | 71 | $1.7 \pm 0.6$ | $0.09 \pm 0.02$ | 28 | $0.19 \pm 0.11$ |
| NC | WT | 84 | $5.9 \pm 0.9$ <sup>*</sup> | $0.56 \pm 0.04$ | 66 | $0.50 \pm 0.08$ <sup>*</sup> |
| NC | γ-T203M | 86 | $4.2 \pm 1.1$ | $0.15 \pm 0.04$ <sup>#</sup> | 40 | $0.44 \pm 0.09$ <sup>\$</sup> |
| NC | β-R196H | 40 | $3.4 \pm 0.9$ <sup>#</sup> | $0.48 \pm 0.05$ <sup>\$</sup> | 23 | $0.54 \pm 0.14$ <sup>\$</sup> |
| NC | 52555#9 + Latr | 37 | $1.1 \pm 1.5$ | $0.13 \pm 0.01$ | 16 | $0.21 \pm 0.08$ |

$n$ , number of cells used in the experiments; $F_{\text{thr}}$ , threshold force required to induce tether formation, was measured at an extraction speed of $4 \mu\text{m s}^{-1}$ ; $k_0$ , quasi-static tether stiffness estimated from the slope of the $F_2$ - $L$ relation; $n_1$ , subset of cells used to determine $\eta_{\text{eff}}$ estimated from the $F_1$ - $v$ relation. Symbols indicate significant differences ( $p < 0.05$ ) of the following comparisons: \* WT NPC vs NC (maturation effect); \$ Latrunculin-A-treated vs mutant cells; # WT vs mutant cells.

**Table 3.** Actin isoform distribution from western blot.

|  | NPC | NC | NPC | NC | NPC | NC |
| --- | --- | --- | --- | --- | --- | --- |
| | WT<br>52555#9 | WT<br>52555#9 | β-R196H | β-R196H | γ-T203M | γ-T203M |
| Actin/GAPDH | $2.7 \pm 0.6$ | $2.1 \pm 0.6$ | $2.3 \pm 0.6$ | $2.3 \pm 0.2$ | $3.6 \pm 0.4$ | $7.2 \pm 0.7$ |
| β-actin | $(46 \pm 11) \%$ | $(63 \pm 17) \%^*$ | $(47 \pm 16) \%$ | $(58 \pm 14) \%$ | $(51 \pm 15) \%$ | $(27 \pm 7) \%^{**}$ |
| γ-actin | $(54 \pm 11) \%$ | $(37 \pm 17) \%^*$ | $(53 \pm 16) \%$ | $(42 \pm 14) \%$ | $(44 \pm 15) \%$ | $(70 \pm 7) \%^{**}$ |
| α-actin | 0 | 0 | 0 | 0 | $(5 \pm 1) \%$ | $(3 \pm 1) \%$ |
| Pan-Actin/<br>GAPDH | $0.8 \pm 0.4$ | $1.1 \pm 0.6$ | $0.8 \pm 0.1$ | $1.2 \pm 0.1$ | $0.9 \pm 0.2$ | $2.2 \pm 0.7$ |

The table reports the results of western blot analysis indicating the % contribution of each isoform ( β-actin, γ-actin, and α-smooth muscle actin) over total actin in WT and mutant cells ( γ-T203M and β-R196H) at the NPC and NC stage and the amount of total actin (quantified with Pan-Actin antibody, see Materials and Methods) expressed as normalized for GAPDH (Pan-Actin/GAPDH). The values are the mean values ( ± SD) obtained from three independent gels for each antibody type. Symbols indicate a significant difference between NPCs and NCs of the same condition (\*p < 0.05; \*\* p < 0.001).

As shown in Fig. 4B, neuronal maturation has the general effect of decreasing *F*_thr_ of all experimental models (Table 2, p always < 0.0001), without significant effects on *k*_0_ (Table 2, p always > 0.4). The dramatic depression of both parameters by Latrunculin-A is the same in NCs as in NPCs.

The *F*_1_-*v* data averaged from WT NPCs of both lines are plotted in Fig. 4C (black circles) and superimposed on those obtained from β-R196H NPCs (blue circles), γ-T203M NPCs (green circles) and Latrunculin-A-treated WT NPCs (magenta circles). Comparison of the linear fits of the relations with Eq. 3 allows to visualize the difference in slope (the viscous coefficient η_eff_) and ordinate intercept (*F*_2’_, the value of *F*_1_ at *v* = 0 for elongations 38-43 μm). The values of η_eff_ estimated by the fit, reported in Table 2, show that the high value of η_eff_ proper of WT NPCs (0.83 pN·s μm^−1^) is slightly and non-significantly reduced, in β-R196H mutants (0.65 pN·s μm^−1^, p = 0.4) while it is significantly decreased in γ-T203M mutants (0.45 pN·s μm^−1^, p < 0.02). The same relations obtained from NCs (Fig. 4D and Table 2) show that η_eff_ decreases upon maturation of WT neurons (0.5 pN·s μm^−1^, p = 0.01) and by a lesser and non-significant extent in β-R196H NCs (0.54 pN·s μm^−1^, p = 0.6), while in γ-T203M NCs it maintains the low value (0.44 pN·s μm^−1^) characterizing the γ-mutant NPCs. An even lower value is observed in Latrunculin-A-treated cells, independent of the degree of maturation (∼0.2 pN·s μm^−1^). The finding that tether elongation in Latrunculin-A-treated cells, irrespective of the degree of differentiation, is characterized by a minimum value of the parameters defining the dynamic (η_eff_) and the quasi-static stiffness (*k*_0_), and the similar finding for *F*_thr_, the threshold force for tether extraction, are straightforward consequences of the primary role of the actin cytoskeleton in cell shape maintenance.

### Effect of neuron geometrical factors on the relevant mechanical parameters

To define the intrinsic mechanical properties of the cell surface, the force responses described above must be scaled for surface-geometry differences across experimental models to obtains the effective force per unit area acting on the cell membrane. This requires considering the size of the surface exposed to the perturbing force, as the area of the patch involved in tether extraction or the dimension of the tether during tether elongation. The tether radius (*r*_t_), measured according to the procedure detailed in Materials and Methods (Fig. S4), is ∼110 nm in WT NPCs and is not significantly affected by either maturation or by β-R196H and γ-T203M mutations (Table S4). In contrast, Latrunculin-A treatment increases the tether radius by a factor of 3. This effect is accounted for by the disruption of the actin cytoskeleton by suppression of both isoform filaments, which by itself reduces the surface tension on the tether (*T*_t_). The relation between *T*_t_ and the force *F* exerted by the tether depends on the tether radius *r*_t_ according to the equation:

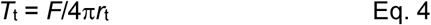

Thus, the reduction in *T*_t_ by Latrunculin-A implies, for a given force, a proportionally larger radius of the tether^28,37–39^.

Since *r*_t_ does not change in the other experimental models (Table S4), except in Latrunculin-A-treated cells, parameters describing tether elongation mechanics, such as *k*_0_ and η_eff_, need not be normalized for tether radius.

Instead, *F*_thr_, the parameter that characterizes the structural changes of the cell surface leading to tether formation, should be normalized for the actual geometry of the cell surface undergoing the mechanical perturbation, that is the area of the membrane patch responding to the initial pulling. Given the limited time resolution of our system, we cannot obtain a direct estimate of the patch area responding to initial pulling for each of the experimental models. On the other hand, considering that membrane-cytoskeleton links are irregularly distributed on the membrane ^26,40^, the patch shape and size interested in membrane-cytoskeleton separation likely change depending on the region of the cell surface, so that a precise patch radius cannot be defined. Under these conditions, we restricted the analysis of tether-formation parameters to raw *F*_thr_ data, considering surface geometry only in relation to soma size.

Through a simple application of Laplace law, limited mainly by the complexity of the soma geometry, one can derive the surface tension *T* of the membrane as 𝑇 ∝ 𝐹 · 𝑟, where *r* is the local radius of curvature of the soma surface.

The reduction of *F*_thr_ in β-mutant NPCs to 8.5 pN (∼0.6 the WT NPCs, Table 2) is the only substantial mechanical difference between WT and β-R196H NPCs. Considering that the soma size, measured from 2D projected images, is reduced to 0.60 (Table 1), its in-plane radius *r* is reduced to 0.77 (see Materials and Methods). From the relation stated above, one can calculate that *T* for the threshold of tether formation (*T*_thr_) of β-R196H NPCs is reduced to (0.6 · 0.77 =) 0.46 that of WT NPCs. Note that if the β-R196H size had remained the same, the mutation-dependent drop of *F*_thr_ in β-R196H NPCs would have been (1/0.77 =) 1.3 times larger than that observed. This suggests that the smaller size of β-R196H NPCs is a compensatory adaptation, as it reduces the vulnerability of the cell surface caused by the mutation-dependent drop in surface tension.

When the same analysis is applied to the effect of the fourfold increase in the soma size (and thus of the twofold increase in *r*) with neuron maturation (Table 1), it comes out that the reduction of *F*_thr_ to 0.4 the NPC value (Table 2) can be accounted for by a maturation-dependent reduction of *T*_thr_ to (0.4 · 2 =) 0.8 the NPC value. Notably, at the NC stage, when the cells have a similar radius and the drop in *F*_thr_ (0.6 the WT value, Table 2) is a direct consequence of the effect of β-R196H mutation on *T*_thr_, the drop in *T*_thr_ is reduced to 0.6 the WT value (it was 0.46 at NPC stage), indicating that the maturation-dependent drop of cell surface resistance mitigates the effect of β-R196H mutation on *T*_thr_.

### β- and γ-actin isoforms are colocalized in the soma and cellular protrusions despite their functional differences

How the alterations of the mechanical responses to external perturbations imposed on the surface of WT and actin mutant neurons at the two stages of maturation are correlated to differences in isoform expression and actin cytoskeleton organization is investigated by western blot and immunocytochemical analyses.

Western blot results (Table 3) show that in wild-type cells, the total actin level, measured as the ratio Actin/GAPDH (see Materials and Methods) and isoform composition remain relatively stable throughout differentiation, with a shift toward β-actin predominance, consistent with existing data^41^. In the β-R196H mutant the actin levels largely overlap the WT profile, indicating minimal impact of this mutation on actin abundance and isoform composition. In contrast, the γ-T203M mutant shows, pronounced total actin upregulation during maturation: the actin/GAPDH ratio doubles between NPC and NC stages, see Materials and Methods for the quantification. The increase is due to specific γ-actin upregulation at the expense of β-actin expression (Table 3, 6^th^ column). Moreover, the γ-T203M mutant displays a small but significant presence of α-smooth muscle actin (α-SMA), a marker that is not typically expressed in neurons.

The evaluation of the relative contribution and localization of each isoform by quantitative confocal immunofluorescence imaging (see Materials and Methods) shows that mutation of one isoform not only causes actin isoform unbalance but it is also associated with specific changes of cytoskeletal architecture. Analysis of reduced-magnification images (20x; Fig. S5) shows that in γ-T203M mutants, β-actin filaments remain predominant, with respect to γ-actin filaments, in contrast to the evidence provided by the western blot (Table 4). This can be explained with γ-T203M mutation implying γ-actin fragmentation by blunting the normal polymerization dynamics. The analysis also confirms the expression of α-SMA in γ-T203M mutants (Fig. S6), in larger proportion in NPCs (26%, Table S5) than in NCs (15%). α-SMA expression likely represents a reaction to the mutation-dependent loss of normal γ-actin cytoskeleton. However, this adaptive response does not compensate for the blunting of the specific function of the γ-actin cytoskeleton.

**Table 4.** β/γ-actin ratio from immunofluorescence imaging.

| Experimental model | | $n$ | $\beta/\gamma$-actin |
| --- | --- | --- | --- |
| NPC | WT | 7 | $1.34 \pm 0.12$ |
| NPC | γ-T203M | 5 | $1.95 \pm 0.15$ |
| NPC | β-R196H | 5 | $1.09 \pm 0.18$ |
| NC | WT | 6 | $1.71 \pm 0.16$ |
| NC | γ-T203M | 5 | $1.28 \pm 0.29$ |
| NC | β-R196H | 5 | $1.29 \pm 0.11$ |

The table reports the $\beta/\gamma$ -actin ratio for WT and mutant cells ( γ-T203M and β-R196H) at NPC and NC stage. Data are presented as mean ± SD, with $n$ indicating the number of images used in the analysis.

The isoform localization by immunofluorescence was optimized by adapting the laser intensity to the isoform-specific fluorophore (Fig. 5, 63x magnification for NPCs (A) and 20x magnification for NCs (B), and Materials and Methods). In all the different experimental models, both in NPCs and NCs (a, WT 52555#9; b, γ-T203M mutant; c, β-R196H mutant) both isoforms are present either in the soma (more markedly at the periphery) or in the protrusions, and their distributions quite clearly overlap, as shown in the third column by merging the images from first and second column.

**Fig. 5.**
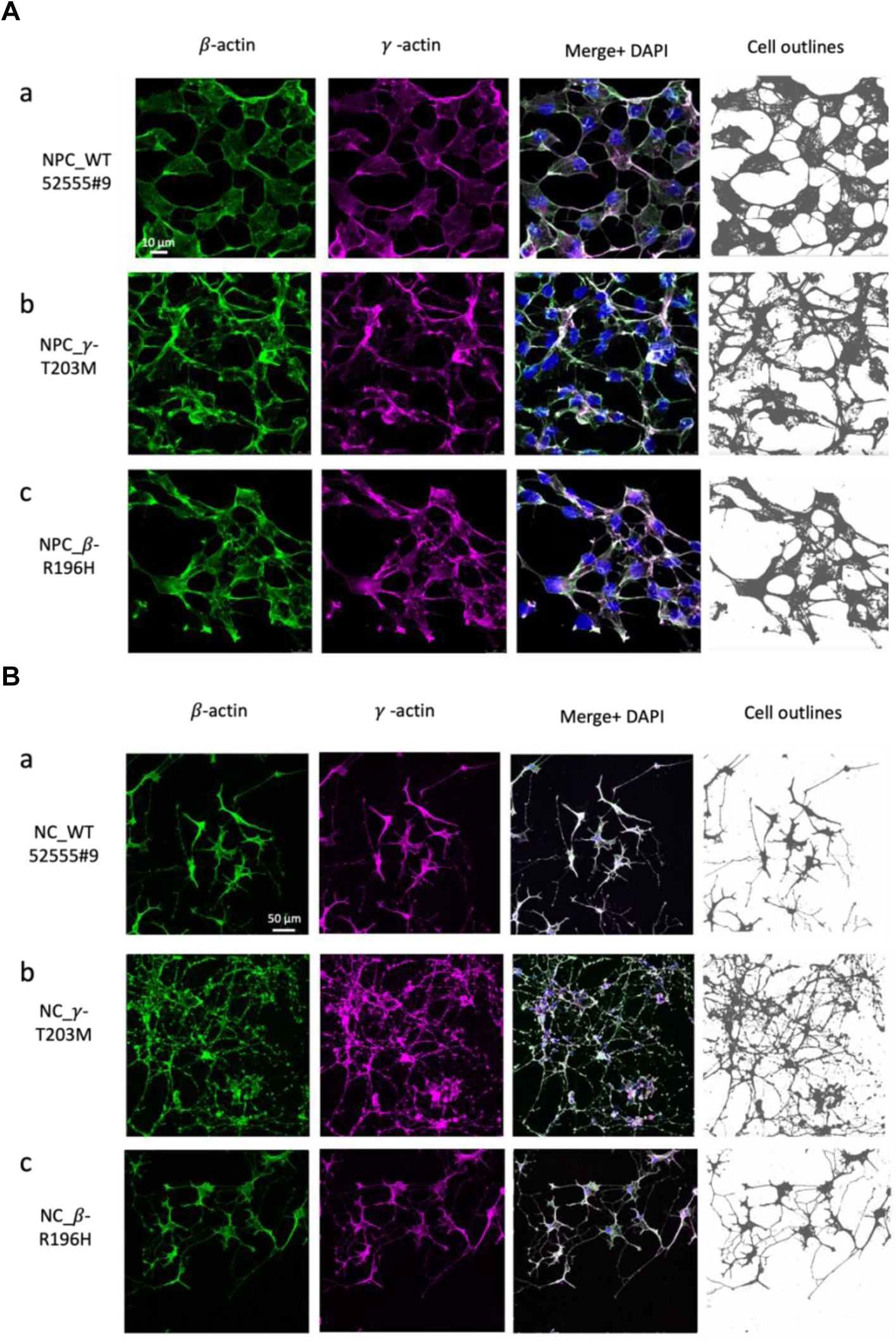
β-actin and γ-actin localization and cell mask differences in WT and mutant cells. **A.** Images of NPCs from WT 52555#9 (a), γ-T203M (b) and β-R196H (c) cell lines. Cells are immunostained for β-actin (green, first column) and γ-actin (magenta, second column); in merged images (third column) white indicates overlap of β- and γ-actin, and nuclei are counterstained with DAPI (blue). Cell mask is maximized in the fourth column (see Methods). Scale bar: 10 µm for NPCs and 50 µm for NCs. **B.** Images of NCs from WT 52555#9 (a), γ-T203M (b) and β-R196H (c) cell lines, immunostained and shown as in A.

Quantitative colocalization analysis confirmed the extensive overlap between γ-actin and β-actin isoforms in both soma and protrusions. Pearson’s correlation coefficients were consistently high across all experimental conditions (0.852-0.959), and overlap coefficients likewise indicated strong spatial coincidence of the two signals (0.895-0.967). When soma and protrusions were analyzed separately, Manders’ coefficients further supported substantial reciprocal overlap between the isoforms (see Table S6).

This finding is not in agreement with previous work ^42^ showing a general strong β-actin enrichment at the cell periphery and a more diffuse distribution of γ-actin within the cytoplasm. Meanwhile, the finding can explain the lack of consensus in the literature on the localization of β- and γ-actin in cells from different tissues (neuronal, epithelial and fibroblast cells)^13,35,43–45^. In this respect, our results support the concept that localization *per se* is insufficient to explain isoform-specific functions; rather, colocalization indicates that the two isoforms have complementary roles, as demonstrated by the distinct consequences of their mutations on the mechanical parameters.

### With neuronal maturation, cell surface resistance decreases as a result of increased cell size and cytoskeletal remodeling

In WT neurons, the resistance to cell surface deformation expressed by the threshold force for tether formation *F*_thr_ becomes 2.5 times smaller upon maturation (Table 2). This can be largely accounted for by the twofold increase in cell radius (Table 1). Consequently, only ∼20% of the decrease in *F*_thr_ is attributable to the maturation-dependent reduction in *T*. However, the limitations of the simplified relation derived from Laplace’s law must be considered not only in relation to the assumption of the soma as a sphere, but also with respect to the contributions of cortical tension and membrane–cortex adhesion/coupling to *T*^46,47^. An important contribution to the maturation-dependent softening is likely to arise from the adaptation of the cytoskeleton–membrane system, which undergoes chemo-structural remodeling, such as a reduction in actin crosslinking and an increase in the turnover of actin–membrane linkers, processes required for neuronal maturation events as neurite outgrowth and synaptogenesis.

Tether elongation serves as a physiologically relevant proxy for the early phases of neuronal connectivity phenomena, such as filopodia protrusions^29^ and intercellular tunneling nanotube connections^30,31^. This idea is supported by the recent time-resolved demonstration of traction-induced tether remodeling with actin recruitment on a timescale of minutes ^27^. With the tether radius remaining constant during neuron maturation (Table S4), the finding that *k*_0_ does not change during maturation (Table 2) is the straightforward demonstration that the underlying property, the *vis a fronte* leading to actin cytoskeleton recruitment within the protrusion, is conserved in the WT neuron independent of its stage of maturation. Pharmacological blunting of actin polymerization with Latrunculin-A almost minimizes *k*_0_ at any stage of WT neuron maturation (Table 2), even if actin is found in the tether after 5 min from tether extraction (Fig. S1). This can be explained if depolymerized actin flows within the tether even faster than polymerized actin, as demonstrated in the tether of fibroblasts treated with the actin polymerization inhibitor cytochalasin D ^27^.

The viscous resistance to tether elongation η_eff_ is halved upon maturation, likely reflecting a maturation-dependent weakening of molecular interactions among cell-surface elements undergoing reciprocal slippage during tether elongation. A substantial contribution could derive from the reduction of membrane-cortex adhesion^46,47^.

### β- and γ-actin mutations differently affect cell surface resistance to perturbations and tether tension

β-R196H and γ-T203M mutations induce distinct effects on mechanical and morphological characteristics of the neurons during differentiation, revealing isoform-specific disruptions in cytoskeletal actin structure-function.

At the NPC stage, β-R196H cells exhibit reduced resistance to cell surface deformation under a force perturbation, marked by the reduction of the threshold force for tether formation *F*_thr_ to 0.6 that of WT NPCs (Table 2). Actually, weakening of cell surface resistance itself is much larger than that shown by the drop in *F*_thr_: when the reduction in soma size of β-R196H mutant is considered, the reduction in tension of the cell surface (*T*_thr_) is 0.46 that of WT NPCs. These considerations, in turn, indicate that the reduction in size of β-R196H NPCs partly compensates for the mutation-dependent loss of cell surface shape stability.

Upon maturation, the resistance of β-R196H NCs against cell surface perturbation, measured by *F*_thr_, is reduced 2.5 times with respect to β-R196H NPCs (Table 2), similar to the reduction of *F*_thr_ upon maturation shown by WT neurons. However, in β-R196H neurons, the soma size is 6 times larger (and thus *r* is 2.45 times larger) in NCs than in NPCs, so that *T*_thr_ = (0.4 · 2.45) remains almost the same. In other words, the whole difference in *F_t_*_hr_ between β-mutant NPCs and NCs is accounted for by the increase in soma size, and maturation does not imply any further change in cell surface deformability beyond that caused by the β-R196H mutation. This contrasts with the maturation-dependent remodeling of the cell surface observed in WT neurons. Thus, as a further consequence of the mutation, β-R196H neurons lose the maturation-associated regulation of cell surface tension.

As far as tether elongation, at the NPC stage, the β-R196H mutation does not significantly impact either of the parameters expressing the viscous (η_eff_) and elastic (*k*_0_) resistance (Table 2). More precisely, η_eff_ of β-R196H NPCs is reduced, but not significantly (−20%, p = 0.4, Table 2) in comparison to that of WT NPCs. Upon differentiation, β-R196H neurons do not show any significant difference in either η_eff_ or *k*_0_ with respect to WT neurons. Accordingly, the actin levels (Table 3) and their isoform distribution in the soma and in the protrusions of both β-R196H mutant NPCs and NCs (Fig. 5, Table S6) largely overlap the WT profile.

γ-T203M NPCs do not show a significant effect of the mutation on the resistance opposed by the cell surface to pulling and tether formation as defined by *F*_thr_ (Table 2). Maturation of γ-T203M mutant induces a drop in the resistance to pulling similar to that shown by WT cells upon maturation and largely, even if not totally, accounted for by the increase in soma size. The most dramatic effects of γ-mutation become evident as far as tether elongation. Both relevant parameters of tether elongation, η_eff_ and *k*_0_, are significantly affected (Table 2). The resistance to slipping between cell surface elements during elongation, marked by η_eff_, is halved in γ-T203M NPCs compared with WT cells, suggesting weakening of molecular interactions between the membrane and cortex, like that observed in WT cells upon maturation. Instead, maturation of γ-T203M cells does not further affect η_eff_, so that the value becomes similar to that of WT NCs. In this respect, η_eff_ appears to be a parameter quite sensitive to regulation, as maturation reduces it to the same value in WT and mutant neurons. Moreover, both β-actin and γ-actin must contribute to WT neuron η_eff_, because the drop caused in this parameter by either β- or γ-T203M mutation is only ∼1/3 of that caused by suppression of both isoform polymerization by Latrunculin-A (Table 2). Ultimately, it should be noted that the small drop of η_eff_ observed in β-R196H NPCs (∼20% and not significant, Table 2) could be due to compensation by the other isoform, partially restoring the molecular interactions underlying membrane-cortex drag.

*k*₀ is reduced by a factor of 4 in γ-mutant NPCs, attaining a level (0.12 pN μm^−1^, Table 2) comparable to that of Latrunculin-A-treated WT NPCs. This minimum value of *k*₀ exhibited by γ-mutant cells is not further affected by maturation and thus remains 4 times smaller than that of WT neurons. The finding that *k*₀ of γ-T203M cells is as low as that of Latrunculin-A-treated WT neurons confirms that the quasi-static stiffness defined by the tether elongation maneuver relies on the recruitment of polymerized γ-actin. Blunting of correct polymerization dynamics by γ-T203M mutation (see also^20,21^) reduces *k*_0_ at any stage of maturation (Table 2), even if actin is found into the tether after 5 min from elongation (Fig. S1). This is because, like in the Latrunculin-A-treated neurons, the resistance opposed to the flow of γ-mutant depolymerized actin within the tether is four times lower than that of polymerized actin.

The functional phenotypes identified by mechanical analysis find their structural correlate in the immunofluorescence imaging of WT and mutant neurons. The images of fixed NPCs and NCs (Fig. 5), labelled with fluorescent antibodies for β-actin (1^st^ column, green) and γ-actin (2^nd^ column, magenta) and merged together with DAPI imaging for nuclei in the third column, demonstrate that in all experimental models β and γ-actin are markedly colocalized. These images can also be exploited for a fine characterization of the soma profile using the corresponding cell masks (4^th^ column), when the colors are reduced to two, grey for the cells and white for the background, and the contrast inside the cell is smoothed (see Materials and Methods). WT NPCs (Fig. 5A) show the edges of the soma that are extended under the tensing effect of straight interneuronal connections. β-R196H NPCs (Fig. 5A-c) substantially maintain this characteristic, while γ-T203M NPCs (Fig. 5A-b) show a fringed soma profile clearly not modelled by interneurons connections that by themselves do not appear tense as they emerge at more acute angle and are randomly curved and fragmented. The quantification of the angle between protrusion direction and the local tangent of the soma boundary (emergence angle θ; Fig. S7A and Materials and Methods) shows that WT NPCs predominantly displayed a small θ (23 ± 19°), indicating that protrusions emerge preferentially with an orientation close to the local soma tangent (Fig. S7B). Cells carrying the β-R196H mutant show a broader distribution of θ shifted towards larger angles (30 ± 18°), which however is not significantly different from that of WT (p = 0.072). The γ-T203M mutant exhibit an even broader distribution further shifted toward a larger angle (40 ± 27°), significantly larger than that of WT. This is a clear consequence of the absence of a tensile action of the protrusion on the soma surface. The difference reflects on distinct morphologies at the NC stage, where WT (Fig. 5B-a) and β-R196H mutant neurons (Fig. 5B-c) show a few well-organized neurites with several filopodia, while γ-T203M mutant neurons (Fig. 5B-b) show numerous fragmented protrusions, without a clear distinction between neurites and filopodia.

The comparative analysis of β-R196H and γ-T203M mutant neurons makes evident that, in contrast to the marked mechanical and morphological differences in γ-T203M mutant, the β-R196H mutant appears quite similar to WT, apart from the specific reduction in cell surface resistance to external stress defined by the drop in *T*. However, a further distinctive element of β-R196H mutation emerges considering the number of cells (*N*_F_) that successfully overcome the fixation protocol for immunofluorescence analysis (Table 5). At the NPC stage β-R196H *N*_F_ is slightly smaller than WT *N*_F_ (−18%, p = 0.6), while it is significantly smaller at the NC stage (−45%, p = 0.05). This finding suggests a larger specific vulnerability of β-R196H NCs that agrees with the reduced resistance of the cell surface to external stress. The vulnerability is less evident at the NPC stage because reduced soma size exerts a compensatory effect.

**Table 5.** Quantification of number of cells overcoming the fixation procedure.

| Experimental models | | $N_F$ | $N$ | $n$ | p |
| --- | --- | --- | --- | --- | --- |
| NPC | WT | $89 \pm 17$ | 4 | 22 | |
| NPC | γ-T203M | $93 \pm 20$ | 2 | 12 | 0.88 |
| NPC | β-R196H | $73 \pm 28$ | 2 | 10 | 0.62 |
| NC | WT | $99 \pm 22$ | 4 | 21 | |
| NC | γ-T203M | $216 \pm 54$ | 2 | 11 | 0.04 |
| NC | β-R196H | $54 \pm 8$ | 2 | 11 | 0.05 |

Total cell number was quantified by counting DAPI-positive nuclei in immunofluorescence images of NPCs and NCs. Data are mean ± SD. $N_F$ indicates the number of cells retained on the coverslip and remaining detectable after the fixation protocol, $N$ represents the number of coverslips considered for each condition, while $n$ indicates the total number of images analyzed.

The *N*_F_ analysis lets a new striking consequence on the blunting of γ-T203M cells maturation to emerge at the NC stage *N*_F_ in γ-T203M is found significantly larger than in WT NCs (2.2 times, p < 0.05), indicating a larger propensity of γ-mutant to proliferate even in the maturing medium. Considering the regulatory effect of tension within a neuron protrusion, this result is expected as a consequence of the fourfold drop of tension within the tether and agrees with the preservation of the morphology of immature neurons.

## Discussion

### Morphological and mechanical clues of the role of the tension within the protrusion of the developing neuron and its blunting by γ-actin mutation T203M

Integrating the morphological characteristics of WT and γ-T203M mutant cells at either stage of maturation with their specific mechanical parameters improves understanding of the mechanisms underlying the development of the WT and γ-T203M mutant phenotypes. At the NPC stage, the high quasi-static stiffness (*k*₀ ∼0.5 pN nm^−1^, Table 2) of the WT cells reflects the high *vis a fronte* driving tether elongation when the ensuing remodeling recruits γ-actin cytoskeleton within the tether. This condition is similar to that naturally occurring during the maturation of a neurite under the tension produced by the growth cone or cell-cell interaction. The corresponding morphological evidence in our experimental model is the tensing effect of interneuronal connections on the profile of WT NPC soma (Fig. 5A-a). The fourfold drop of *k*₀ in γ-T203M NPCs (Table 2) is matched to the loss of the tensing effect of interneuronal connections on the soma profile (Fig. 5A-b) quantitatively estimated by the increase in the mean emergence angle of the protrusion (Fig. S7B). The consequences on the failure of the maturation process are evident in the last column of Fig. 5B. The comparison of γ-T203M NCs (b) to WT NCs (a) clearly demonstrates that the γ-mutant neurons are characterized by a diffuse network of connections missing the transition to a connectivity made by few selected neurites.

The conclusion above reveals that axon extension and connectivity in differentiating neuronal networks are based on mechanotransduction processes underpinned by traction forces exerted directly by the neurite. This requires a correct polymerization dynamics of γ-actin, allowing the mechanical coupling between the soma and the periphery explored by the neurite. In our experiment, the parameter that allows defining the correct γ-actin polymerization as the condition for neuron maturation is the quasi-static stiffness *k*_0_. In fact, the development of a correct connectivity is prevented in γ-T203M neurons by a drop in *k*_0_ that is similar to that produced in WT neurons by the actin polymerization inhibitor Latrunculin-A.

### The β-actin mutation R196H reduces cell surface resistance to external perturbations, thereby increasing neuronal vulnerability

A key finding from the parallel analysis of β-R196H mutant cells is that mechanical and imaging readouts closely resemble those of WT cells, prompting the question of which functional deficit accounts for the pathogenicity associated with the β-R196H mutation. The only significant differences between the β-R196H mutant and the WT phenotypes are, on the one hand, the reduction in size of the β-R196H NPCs (Table 1) and, on the other hand, the reduction of *F*_thr_ (Table 2). Reduction of *F*_thr_ does not appear to be a striking effect of the mutation when compared to that reflecting the regulatory effect of WT neuron maturation, but a striking difference emerges when the surface tension is considered: *T*_thr_ in β-R196H NPCs is reduced to 0.46 of the value in WT NPCs and does not show any further change upon maturation. A further distinctive element for β-R196H mutant pathogenicity appears the intrinsic vulnerability of β-R196H NCs, suggested by the reduced number of cells (*N*_F_) that successfully overcome the fixation protocol for immunofluorescence (Table 5). The finding that, at the NPC stage, β-R196H *N*_F_ is not significantly smaller than WT *N*_F_ is a further demonstration of the compensatory effect of the reduced size of β-R196H NPCs further supports the compensatory effect of reduced β-R196H NPC size. The *N*_F_ analysis shows a larger propensity of γ-T203M mutant to proliferate even at the NC stage. This agrees with the conclusions of both mechanical and imaging analyses. The loss of tension within the tether in γ-actin mutant reproduces the conditions that, based on the evidence reported in literature ^5,6^, prevent progression to the differentiation phase that normally inhibits proliferation and suppresses multiple protrusions.

### *In situ* Identification of distinct isoform-dependent functions of actin cytoskeleton is a prerequisite for the detailed description of the underlying molecular mechanisms

The mechanical parameters *T*_thr_ and η_eff_ appear to share a common molecular basis in the interaction between the actin cortex and the proteins constituting the membrane-cortex adhesion system. A contribution to future investigation on the detailed description of the distinct underlying mechanisms is the demonstration here that the two parameters are differently affected by either actin isoform mutation: *T*_thr_ is specifically decreased by β-R196H mutation and unaffected by γ-T203M mutation, while η_eff_ is more affected by γ-T203M mutation and less by β-R196H mutation. In β-R196H NPCs, γ-actin may complement β-actin, partially restoring normal behavior. These conclusions contribute to the clarification of the question that, even if the filaments of the two isoforms are colocalized in the cortex (Fig. 5; Table S6) this spatial overlap does not imply mechanical redundancy, because force transmission depends on filament dynamics, filament organization, and protein interactions rather than localization alone. Thus, β- and γ-actin networks likely provide non-equivalent mechanical contributions within the developing neuronal cortex ^13–15^, as indicated by the different effects of their mutations: *T*_thr_ is affected selectively by β-R196H mutation, suggesting that β-actin filaments contribute to membrane-cortex resistance against shape change induced by a stress on the cell surface, while η_eff_ is affected to a higher degree by γ-T203M mutation, suggesting that γ-actin filaments contribute to molecular interactions opposing reciprocal slippage between elements in the cell surface during tether elongation.

## Materials and Methods

### Ethics approval and sample handling

Clinical data and dermal samples were collected at TU Dresden from individuals carrying *ACTB* or *ACTG1* variants. Recruitment and sampling were approved by the Ethics Committee of TU Dresden (EK 127032017 and EK 44012019), and all participants provided written informed consent; longitudinal clinical data were recorded in REDCap. For experiments performed at the University of Florence, human-derived samples were received fully anonymized, identified only by an alphanumeric code, in according with GDPR (UE 2016/679), used exclusively for mechanical and biophysical analyses (no genetic testing), and not stored beyond the experimental period. Additional ethical clearance was granted by the CNR *Commissione per l’Etica e l’Integrità nella Ricerca* (26 April 2021).

### Cell lines and experimental models

Human iPSC (hiPSC) lines were obtained from four female donors: two patients with BWCFF syndrome and two WT controls. The two patient-derived hiPSC lines carried either *ACTB* mutation p.R196H (49113#16) or *ACTG1* mutation p.T203M (32377Da9). A first WT line used for the experiments was derived from donor dermal fibroblasts (TU Dresden, 52555#9), while a second was a commercially available healthy control line (SCTi003-A; STEMCELL Technologies). Preliminary morphometric and mechanical analyses demonstrated that the two WT lines did not differ in any of the measured morphological or mechanical parameters (Table S1 and S3). In addition, immunocytochemistry/confocal imaging (both qualitative morphology and quantitative β/γ-actin fluorescence readouts) did not reveal detectable differences between the two WT lines. Therefore, data obtained from the two WT lines were pooled and collectively referred to as WT throughout. Cells were analyzed at the neural progenitor cell (NPC) stage (corresponding to 18 to 25 days after the start of neural induction) and as mature neurons (NC) (corresponding to days 47 to 50 after hiPSC seeding, i.e., 14 days after transfer to neuronal maturation medium). See also Fig. 1 for further details of the timeline.

### hiPSC reprogramming, neural induction and differentiation

Reprogramming was performed at TU Dresden using the CytoTune-iPS 2.0 Sendai Reprogramming Kit (^48^ Thermo Fisher Scientific), delivering non-integrating Sendai vectors encoding OCT4, SOX2, KLF4 and c-MYC, starting from donor-derived dermal fibroblasts. Karyotype analysis was performed in-house for all donor-derived hiPSC lines, whereas the commercially obtained WT hiPSC line was provided with supplier-certified karyotype documentation. Only lines with normal karyotype and negative for mycoplasma and residual Sendai virus were used. hiPSCs were maintained on Matrigel-coated plates in mTeSR1 medium (STEMCELL Technologies) at 37 °C and 5% CO₂ with daily medium changes, passaged at 80-90% confluency using enzyme-free reagent used to selectively detach and passage hiPSCs (ReLeSR), and cryopreserved in serum-free cryopreservation (freezing) medium (mFreSR) using a controlled-rate freezing protocol. Neural induction was performed with the STEMdiff Neural Induction kit ^49^ by seeding dissociated hiPSCs into ultra-low attachment 96-well plates in a Neural Induction Medium (STEMdiff) supplemented with SMADI inhibition and ROCK inhibitor to form embryoid bodies, which were transferred to Matrigel-coated plates to generate neural rosettes; rosettes were manually selected and expanded into NPCs maintained from passage 3 onward in Neural Progenitor Medium (STEMdiff). NPCs were subsequently differentiated into neurons using the STEMdiff Forebrain Neuron Differentiation and Maturation Kits ^50^, maintained in differentiation medium for 5-7 days and then switched to maturation medium. From day 14 onward, NC were analyzed as the standard time point for downstream analyses.

### Characterization of cell morphology, maturation and actin expression

#### Bright-field morphometry

Bright-field microscopy was used to monitor morphology as a qualitative readout of differentiation/maturation and to quantify cell size and shape in WT and mutant models at defined stages (NPCs and NCs). Images were acquired with a Nikon Eclipse Ts2 inverted microscope using identical settings within each experiment (NPCs: 20× air, NA 0.75; NCs: 10× air, NA 0.25). Non-overlapping fields were collected while avoiding crowded areas to enable reliable single-cell segmentation. Images were analyzed in Fiji/ImageJ ^33^ using a standardized pipeline (8-bit conversion, background correction when needed, thresholding, mask refinement, and removal of debris/aggregates). Only isolated cells with clear boundaries were included. Cell area, perimeter and circularity were extracted with the Analyze Particles tool using identical criteria across conditions.

### Immunofluorescence protocol, confocal microscopy and quantitative image analysis

Immunofluorescence analyses were performed on NPCs and NCs to evaluate neural identity, β-and γ-actin expression, actin cytoskeletal organization and the proportion of α-smooth muscle actin (α-SMA) positive-cells. 1×10^4^ cells were plated into individual wells of a 24-well plate and allowed to adhere to 12-mm glass coverslips previously coated with Matrigel and maintained in a humidified cell culture incubator under standard culture conditions (37°C and 5% CO₂). NPCs were fixed 24 h after plating, whereas NCs were fixed 14 days after transfer to maturation medium. Cells were fixed with 1% paraformaldehyde for 30 min at room temperature (RT), followed by permeabilization with cold methanol (−20 °C) for 5 min. After rehydration in phosphate-buffered saline (PBS), cells were blocked with 2% bovine serum albumin (BSA) for 1 hour at RT. Samples were incubated with primary antibodies for 1 hour at RT, followed by fluorescent secondary antibodies together with DAPI for 30 min at RT in the dark. After PBS washes, coverslips were mounted in Mowiol. Antibody sources, catalog numbers, and working dilutions are listed in Tables S7 and S8.

Neural identity was confirmed by immunostaining for SOX2 and Nestin in NPCs and Tuj1 in NCs (Fig. S8).

Confocal imaging was performed using a Leica SP8 CLSM (Mannheim, Germany) with a 63x oil objective (HC PL APO CS2 63x/1.40 OIL) for NPCs and a 20x dry objective (HC PL APO CS2 20x/0.75 DRY) for NCs, unless otherwise specified. For each sample, z-stacks were acquired at 2048 × 2048 pixels with a scan speed of 200 Hz and a Z-step size of 1 μm. Acquisition parameters, including laser power, detector gain, and photomultiplier settings ^51^, were kept constant across all experimental models and genotypes. Care was taken to avoid signal saturation in any plane, ensuring accurate quantitative representation of fluorescence intensity. Maximum intensity Z-projections were subsequently generated from the z-stacks for further analysis.

For semi-quantitative fluorescence analysis of β-actin and γ-actin (Fig. S5), channels were separated, and background fluorescence was subtracted. Mean fluorescence intensity for β-actin and γ-actin was measured on the projected images for each field, averaged per coverslip, and then averaged across biological replicates. The β/γ-actin ratio was calculated from the corresponding mean intensities. Analyses were performed blinded to genotype using at least five non-overlapping fields per condition.

α-Smooth muscle actin (α-SMA) immunostaining was performed to assess the proportion of cells expressing this isoform in each experimental model (Fig. S6). α-SMA-positive cells were identified on projected images and expressed as the percentage of total DAPI-positive nuclei per field (Table S5).

Qualitative assessment of soma contour, protrusion morphology, and actin organization was performed on high-magnification confocal images (Fig. 5). For this purpose, maximum-intensity projections were generated, and cell masks were produced in Fiji/ImageJ using a semi-automatic segmentation workflow based on contrast enhancement and intensity thresholding, followed by manual refinement when necessary to accurately delineate cell boundaries. For visualization, masks were rendered with colors reduced to grey for the cell and white for the background.

Colocalization was quantified using the JaCoP plugin^52^. For each sample, two images were analyzed by selecting from each z-stack five planes for NPCs and six planes for NCs (corresponding to 5 µm and 6 µm, respectively). Pearson’s correlation coefficient (PCC) was used to assess the linear correlation between pixel intensities, whereas Manders’ coefficients (M1 and M2) measured the fraction of signal overlap between channels. Values were calculated on the selected planes, excluding the DAPI channel, to evaluate the degree of protein co-distribution. Higher coefficient values indicate stronger colocalization between the signals (see Table S6).

### Quantification of protrusion emergence angle

To quantify the orientation of cellular protrusions relative to the soma boundary, a geometrical method was implemented using Fiji/ImageJ, as detailed in Fig. S7. The contour of the cell body (soma) was first approximated by fitting an ellipse to the visible boundary of the soma (Fig. S7A, red contour). This approximation provides a smoothed representation of the global soma shape and reduces the influence of local membrane irregularities that could affect the estimation of the local boundary orientation. Similar centroid-based geometrical descriptions are commonly used in quantitative analyses of cell edge dynamics and protrusive activity ^53,54^. This analysis provides a robust estimate of the local membrane orientation and is consistent with approaches used to quantify protrusive activity relative to the cell edge ^55^. The protrusion emergence angle θ is measured relative to the local tangent of the soma boundary as:

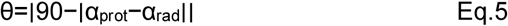

where α_rad_ is the orientation of the radial vector (yellow continuous line in Fig S6A and α_prot_ is the orientation of the protrusion axis (green line). The protrusion emergence angle therefore represents the deviation of the protrusion axis from the local tangent of the soma boundary. Because protrusion orientation is an angular variable and preliminary inspection of the distributions indicated deviations from normality, comparisons among the three experimental groups were performed using the non-parametric Kruskal-Wallis test ^56^. Pairwise comparisons were then carried out using two-sided Mann-Whitney U tests ^57^, and uncorrected p-values are reported. To further assess orientation bias, protrusion emergence angles were grouped into 20° bins (0-20, 20-40°, 40-60°, and 60-90°), based on the distributions shown in the polar plots (Fig. S7B), and the frequency distribution across conditions was compared using a chi-square test of independence.

### Quantification of cells passing fixation

To quantify the number of cells after fixation and staining, DAPI-positive nuclei were automatically segmented and counted in Fiji/ImageJ using thresholding and the Analyze Particles tool. Counts were performed on all acquired fields for each coverslip, and the number of nuclei per image was used for statistical comparison among experimental conditions. *N*_F_ was defined as the number of cells retained on the coverslip and detectable after fixation and immunofluorescence staining. This parameter was used as an operational readout of cell persistence through the fixation/staining workflow and, in mechanically fragile conditions, may reflect increased vulnerability to experimental handling. Statistical comparisons of *N*_F_ values were performed using two-way ANOVA (p < 0.05).

### Western blot analysis

Western blot analysis was employed to support the quantification of actin isoforms expression. For this, cells were lysed in RIPA buffer supplemented with sodium orthovanadate, protease inhibitor cocktail, and PMSF. Total protein content was quantified using the Qubit Protein Assay (Thermo Fisher Scientific), and equal amounts of protein (10 μg) were run on NuPAGE Bis-Tris gels and transferred to nitrocellulose membranes using iBlot 3. Membranes were blocked in 5% non-fat milk in Tris-buffered saline + Triton x-100 (TBS-T), incubated with primary antibodies and HRP-conjugated secondary antibodies (antibody sources, catalog numbers and working dilutions are listed in Tables S9 and S10), and developed using SuperSignal West Pico PLUS; chemiluminescent signals were acquired with a ChemiDoc system (Bio-Rad). Band intensities were quantified in Fiji /ImageJ, normalized to GAPDH, and analyzed from at least three independent biological replicates per condition. The content of β-, γ-, and α-smooth actin was expressed as relative abundance with respect to the sum of all isoform values. Total actin was expressed as this sum normalized for GAPDH (Actin/GAPDH) or directly quantified over GAPDH with an antibody designed to detect total actin by recognizing epitopes common to all actin isoforms (Pan-Actin/GAPDH, see Table 3). Statistical analyses were performed using two-way ANOVA (p < 0.05).

### Mechanical experiments

To characterize cell surface mechanics a custom-built Dual-Laser Optical Tweezers (DLOT) with high-resolution and high dynamic range in force and movement ^24,58,59^ is used. The DLOT consists of two counter-propagating 808 nm diode lasers (250 mW; Lumics GmbH) with orthogonal polarizations focused through opposing water-immersion objectives (60×, NA 1.20; Olympus) to generate a stable optical trap. This configuration enables force measurements up to 250 pN, with ∼0.3 pN resolution. Force detection is based on position-sensitive detectors that exploit the conservation of light momentum ^60^. Temperature was maintained at 37 °C with fluid-based thermal regulation ^61,62^. Bright-field images of bead-cell interaction were acquired with a CCD camera (Bitron Group spa) under transmitted illumination (470 nm LED). Instrument control and acquisition were managed through custom LabVIEW software (National Instruments Corporation) with sampling rates up to 100 kS/s. NH₂-functionalized polystyrene beads (radius 1.5 μm; Kisker Biotech GmbH & Co. KG) were suspended in HBSS and introduced into Matrigel-coated chambers containing adherent cells mounted on a three-axis nanomanipulator stage (nano-PDQ375, Mad City Labs Inc). The chamber allowed rapid solution exchange. Experiments were performed in Hanks’ Balanced Salt Solution (HBSS) at physiological pH at 37°C, under CO₂-free conditions. The cell soma was brought into contact with the trapped bead (initial, or 0, position), selecting a region of the soma surface at least 3 μm away from the nucleus to avoid mechanical bias from the stiffer perinuclear region ^63^. To promote stable bead-membrane adhesion, the cell was translated toward the bead along the cell adhesion plane until a small membrane indentation was observed, which was maintained for at least 10 s. Then tether extraction/elongation was performed by translating the microscope stage away from the trapped bead using a ramp-shaped displacement at a velocity ranging from 1 to 8 μm/s, with an amplitude ranging from 10 to 50 μm. At the end of each ramp, the bead-cell distance was kept constant (hold phase) for 10-30 s, after which the cell was returned to the initial position. A pause l of 15-100 s was introduced between consecutive pulling cycles. Details on the velocity and amplitude of the elongation, duration of the hold, and total period of each cycle are reported in Results. Force responses to length perturbations were analyzed using Igor Pro 8 and custom LabVIEW scripts. Statistical significance (p < 0.05) was assessed using two-way ANOVA, and data are presented as mean ± standard deviation. For each cell, multiple tether elongation-relaxation cycles were performed, and recordings were excluded in the presence of unstable bead trapping or bead loss, failure to form a stable tether, visible cell detachment from the substrate, excessive stage drift or focus loss during the ramp or hold phases, or evident changes in cell shape. Force responses and derived mechanical parameters (*F*_thr_, *k*_0_ and η_eff_) were extracted from each cell, that was treated as a single independent experimental unit.

#### DLOT bright-field imaging and tether geometry measurements

Bright-field DLOT imaging was used to visualize tether formation and to quantify tether radius as a measure of membrane–cortex coupling. All quantitative analyses of tether geometry in WT cells were performed on the pooled dataset obtained from the two WT lines, as no significant differences in tether radius or image-based morphological descriptors were detected between the two WT lines. Calibration in the x-y plane was performed by applying controlled nanopositioner displacements and measuring the corresponding image shifts. Tether images were processed in Fiji/ImageJ using background subtraction, thresholding, binary mask generation, and averaging over multiple frames. Only sharply focused tether segments were analyzed. Because the tether width approached the diffraction limit, the tether radius was estimated after correction for optical resolution. The system point spread function (PSF) was measured using 100 nm fluorescent beads (TetraSpeck; Invitrogen). Gaussian fits yielded σ_A_, corresponding to the PSF width (instrumental broadening), and σ_o_, corresponding to the observed width of the tether intensity profile. The deconvolved tether width was then calculated as σ_t_ = √ (σ_o_² - σ_A_²), and tether radius was approximated as *r*_t_ ≈ 2σ_t_. *r*_t_ was operationally defined from the deconvolved Gaussian width as described above. Representative tether images and the corresponding Gaussian intensity-profile fits are shown in Fig. S4, whereas the resulting *r*_t_ values are reported in Table S4.

#### Live-cell fluorescence imaging of actin in extracted tethers

To visualize the presence of actin within extracted membrane tethers, all experimental models (WT, mutant, and Latrunculin-A-treated cells) were stained live with CellMask™ Orange Actin Tracking Stain (Thermo Fisher Scientific). Live-cell fluorescence imaging was performed on a Zeiss Axioscope 7 microscope equipped with a 63× oil-immersion objective (NA 1.40), using a Zeiss Colibri fluorescence system for excitation at ∼545 nm and emission collection at ∼570 nm. Membrane tethers were mechanically extracted using the DLOT system and subsequently immobilized on Matrigel-coated glass coverslips within the experimental chamber by switching off one laser and using the second to push the bead onto the coverslip surface. The chamber was then removed from the stage and promptly transferred to the Zeiss microscope for live fluorescence imaging, with image acquisition initiated within approximately 5 min after tether extraction (see Fig. S1).

#### Latrunculin-A treatment

To evaluate the specific contribution of actin filaments to cell mechanical properties WT cells were treated with 5 µM Latrunculin-A (Merck) for 20 min at 37 °C in a cell culture incubator. Latrunculin-A, a marine toxin that binds monomeric (G-)actin and prevents filament assembly, was used to induce actin depolymerization and thereby test the effects of transient disruption of the actin network. Following treatment, cells were washed with pre-warmed medium and and imaged by live-cell fluorescence microscopy after staining with CellMask™ Red Actin (Thermo Fisher Scientific) to visualize cortical and filamentous actin structures. The observed loss of F-actin organization confirmed effective depolymerization (see Fig. S1).

## Acknowledgments

The authors thank the members of the PredACTINg Consortium for helpful discussions. The technical contribution of Vera Schäffer from the Department of Human Genetics, Hannover Medical School, for assistance with ultracentrifugation and the Western blot workflow, is gratefully acknowledged. The authors also thank Prof. Gabriella Piazzesi, Prof. Sandra Citi, and Prof. Fabio Benfenati for their careful reading of the manuscript and for their valuable comments, which contributed to improving the paper. This work was supported by the European Joint Programme on Rare Diseases 2019 (PredACTINg, EJPRD19-033), funded by the European Union’s Horizon 2020 research and innovation programme under Grant Agreement No. 825575, with support from the Italian Ministry of University and Research (DM 1638), and by the Italian Ministry of University and Research (PRIN 2022, MAS-NeurActin, 2022XJ29R7). JG was supported by the PREPARE program for medical scientists from Hannover Medical School.

## Author Contributions

EB and PB designed the study. Mechanical experiments were performed by EB, IP, and PB. Western blot and cell growth analyses were carried out by EB, JG, and IN. Data analysis was performed by EB, IP, MR, and PB. Integrated mechanical, morphological, and fluorescence results were interpreted by VL, MR, IP, EB, and PB. Confocal imaging and image analysis were performed by EB, GM, and CDA. NDD provided the cell lines. The manuscript was written by PB and VL, with contributions to drafting and revision from MR, IP, EB, NDD. All authors reviewed the manuscript and approved the final version.

