## Supporting Information for "The distinct role of actin isoforms on mechanosensing-based maturation of hiPSC-derived neurons unveiled by isoform-specific mutations"

**This PDF file includes:**

Figures S1 to S8
Tables S1 to S10

**Figure S1.**

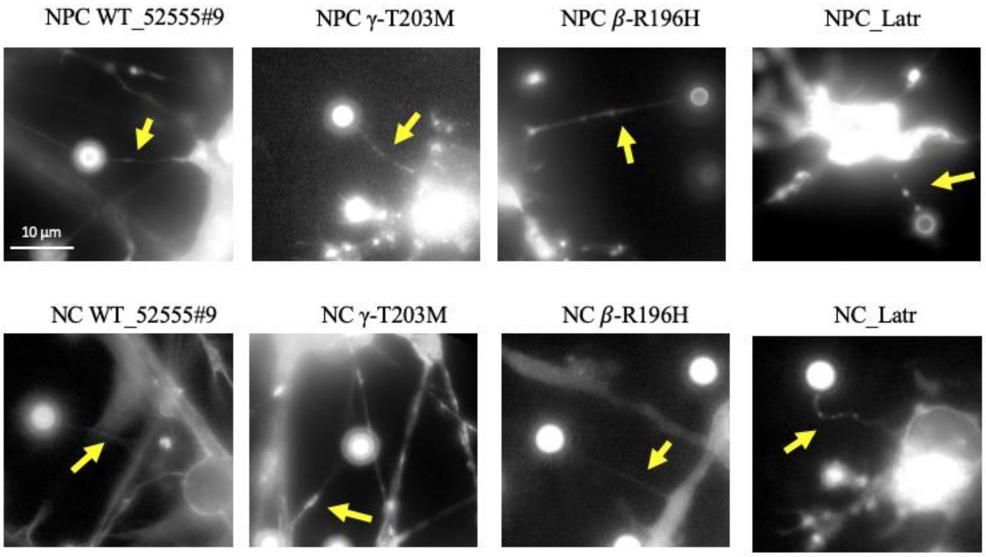

**Fig. S1. Presence of actin in mechanically extracted tethers.**
Representative fluorescence images of tethers extracted from NPCs and NCs for all experimental models. Cells were preloaded with CellMask™ Orange to visualize actin prior to DLOT experiments (see Materials and Methods). Yellow arrows indicate actin within the tether, visible as thin fluorescent structures extending from the plasma membrane to the bead. In control and β-mutant cells, actin within the tether appears continuous and structurally preserved. In γ-mutant cells and in cells treated with Latrunculin-A, actin fluorescence along the tether appears discontinuous or fragmented. Scale bar: 10 µm.

**Figure S2.**

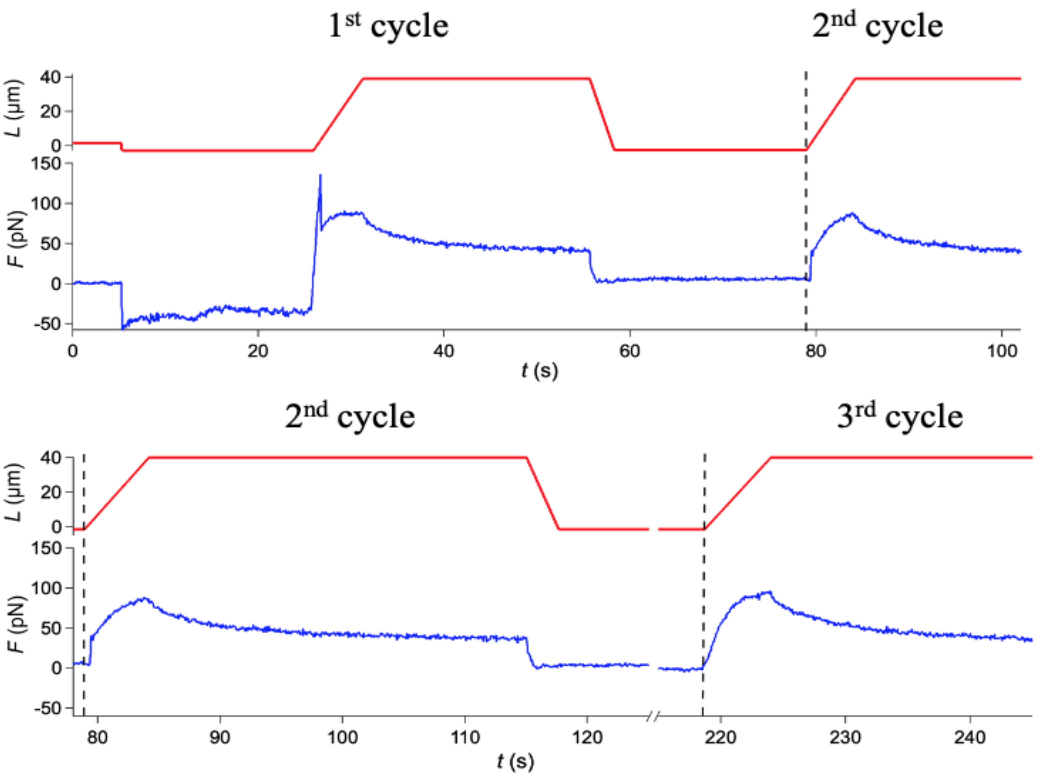

**Fig. S2. Mechanical characteristics of tether extraction and elongation in three consecutive cycles separated by different hold time durations.**
Force response (*F*, blue) to the displacement (*L*, red) imposed on a WT NPC (the same as Fig.1). Tether extraction is achieved by moving the cell away with a ramp-shaped elongation (velocity 8 µm $s^{-1}$, amplitude 45 µm). At the end of the ramp, the position is held for ~30 s, after which the cell is returned to the initial position. The cycle is repeated after a pause of: ~25 s between 1st and 2nd and ~105 s between 2nd and 3rd cycle. Vertical dashed lines mark the start of 2nd and 3rd elongation phase. Following the ~ 25 s pause, the start of force rise lags (0.6 s) the start of elongation. The shorter delay with respect to that at the start of the second cycle in Fig. 2 is accounted for by the twice faster elongation speed. The 3rd cycle elongation, starting after a pause of 105 ms, induces an almost simultaneous increase in force.

**Figure S3.**

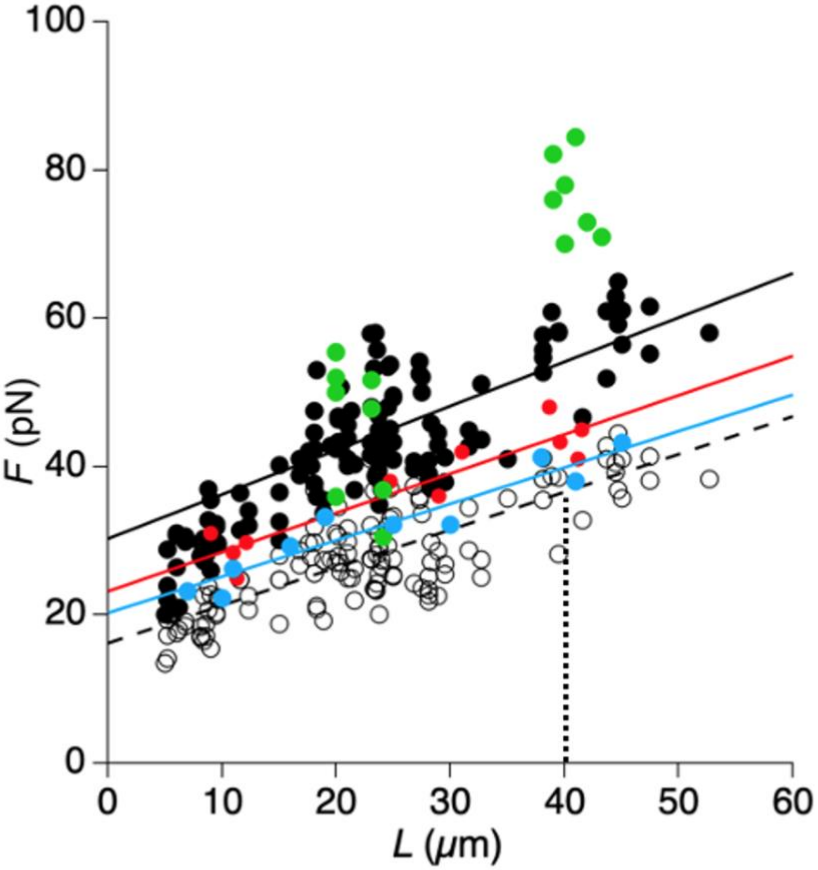

**Fig. S3. Dependence of $F_1$ and $F_2$ on tether elongation at different velocities.** Filled circles are the $F_1$ values measured at different elongation velocities: 1 μm s$^{-1}$ (light blue), 2 μm s$^{-1}$ (red), 4 μm s$^{-1}$ (black), and 8 μm s$^{-1}$ (green). Solid lines represent linear fits to the $F_1$-$L$ data points at each velocity for elongation time satisfying the condition $L/v > t_r$ (the risetime of the viscoelastic component). Open circles are the $F_2$ values from the same records as $F_1$. As expected from Fig.3B, $F_2$ points contribute to a unique relation independent of $v$. The black dashed line represents the linear fit to the $F_2$-$L$ data. For each elongation velocity tested in the range 1–8 μm s$^{-1}$, the slopes of the $F_1$-$L$ relations obtained at different velocities estimate the quasi-static elastic stiffness $k_0$. The finding that they are quite similar demonstrates their independence of $v$. In WT NPCs, $t_r$ = 4.4 s; therefore, at $v$ = 8 μm s$^{-1}$ (the highest elongation velocity tested) the minimum elongation required to exceed $t_r$ was $L = v \times t_r$ = 8 μm s$^{-1}$ × 4.4 s ≈ 35 μm. Since the maximum elongation attainable with the experimental apparatus was approximately 40 μm, at 8 μm s$^{-1}$. only the largest elongations fulfilled the requirement. As a consequence, an independent linear fit of the $F_1$-$L$ relationship could not be reliably achieved for this velocity, the available valid data being clustered around $L$ ≈ 40 μm. Nevertheless, these $F_1$ values were included in the subsequent $F_1$-$v$ analysis (Fig. 3E), as they lay on the extrapolation of the linear relations defined by the lower-velocity data. For elongations reached at times > $t_r$, specifically in the range of approximately 38-43 μm, $F_1$ increased linearly with $v$. The resulting $F_1$-$v$ relation (Fig. 3E) was fitted with Eq. 3 to estimate the effective viscous resistance $\eta_{eff}$.

**Figure S4.**

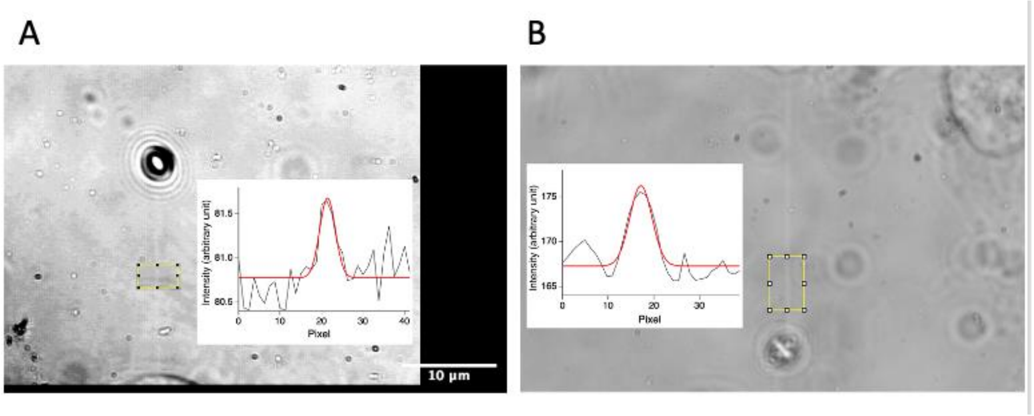

**Fig. S4. Tether size measurements by bright-field DLOT imaging.** Tether images from a WT NPC (A) and a Latrunculin-A-treated NPC (B). The insets show the transverse intensity profiles measured across the tether within the yellow rectangles, with the Gaussian fits used to estimate tether width superimposed on the corresponding profiles. To avoid artifacts due to shadowing or out-of-focus effects, the tether diameter was quantified exclusively within the sharply focused portion of the tether image, corresponding to the boxed region. Scale bar: 10 μm.

**Figure S5.**

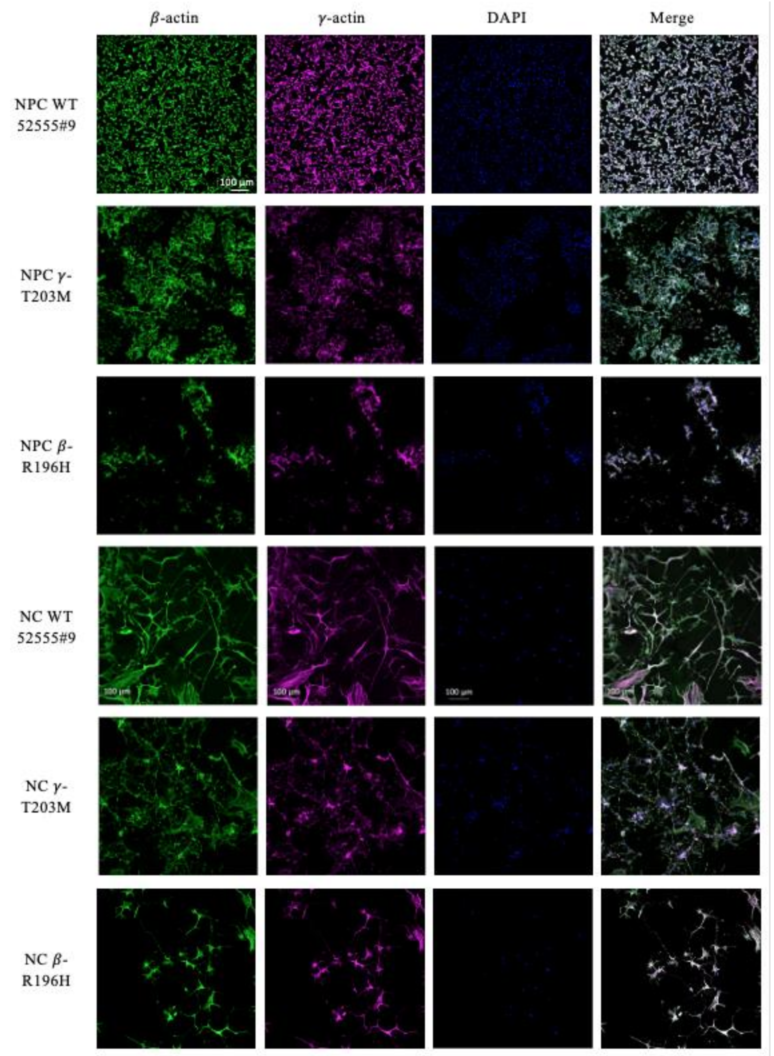

**Fig. S5. Immunofluorescence images used for quantitative analysis of β- and γ-actin distribution.** Confocal images of NPCs and NCs derived from WT (52555#9), γ-T203M and β-R196H cell lines. Cells were stained for β-actin (green), γ-actin (magenta) and nuclei (DAPI, blue); merged images are shown in the right column. Scale bar: 100 µm.

**Figure S6.**

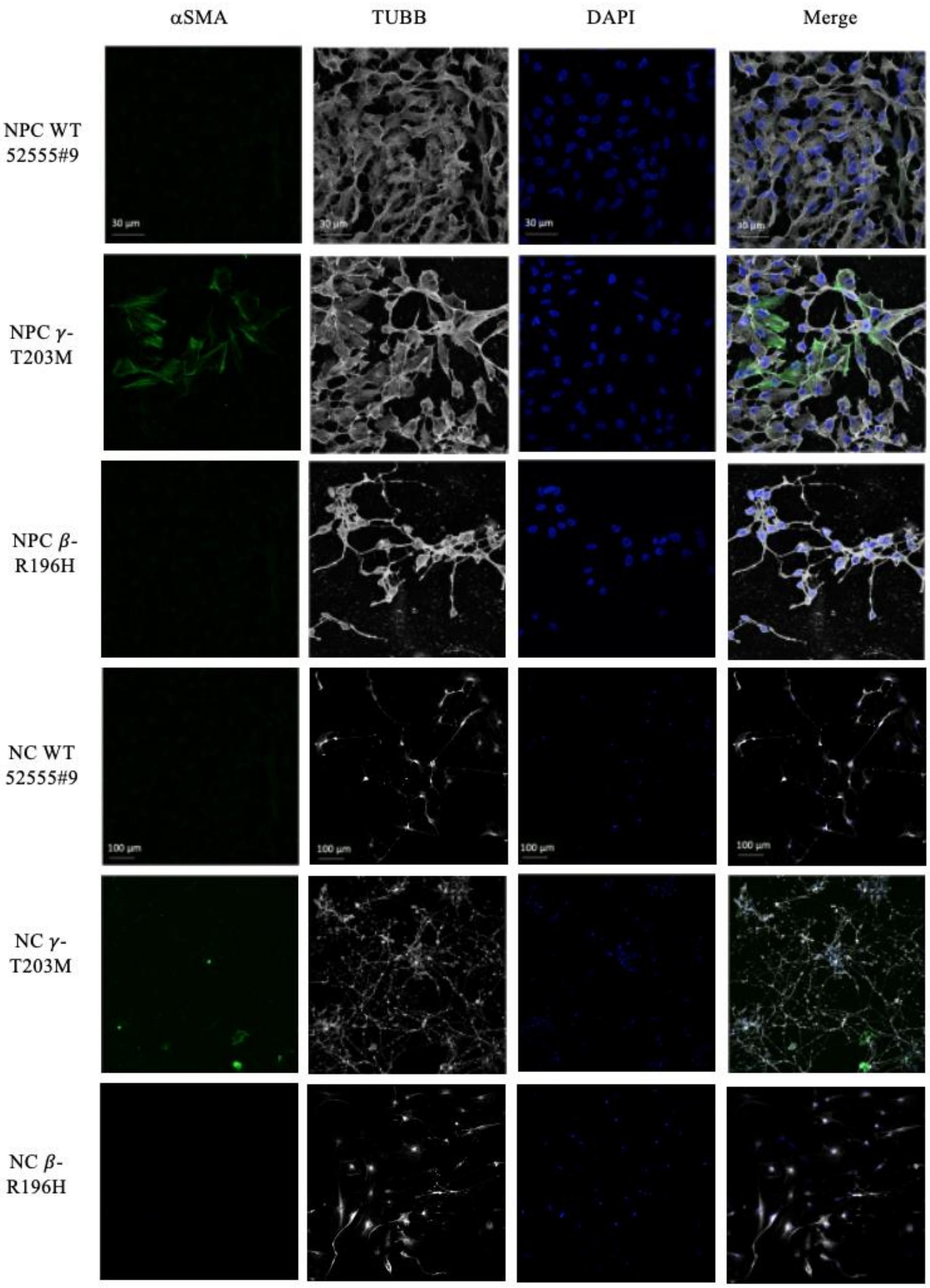

**Fig. S6. Expression of α-smooth-muscle actin and TUBB in WT and mutant both NPCs and NCs.** Images of NPCs and NCs derived from WT (52555#9), γ-actin T203M and β-R196H cell lines. Cells were stained for α-smooth muscle actin (α-SMA, green), β-tubulin (TUBB, gray), and nuclei (DAPI, blue). The merged images illustrate the co-localization of cytoskeletal markers with nuclear staining. Images were acquired as detailed in Materials and Methods. Scale bars: 30 μm (NPCs) and 100 μm (NCs).

**Figure S7.**

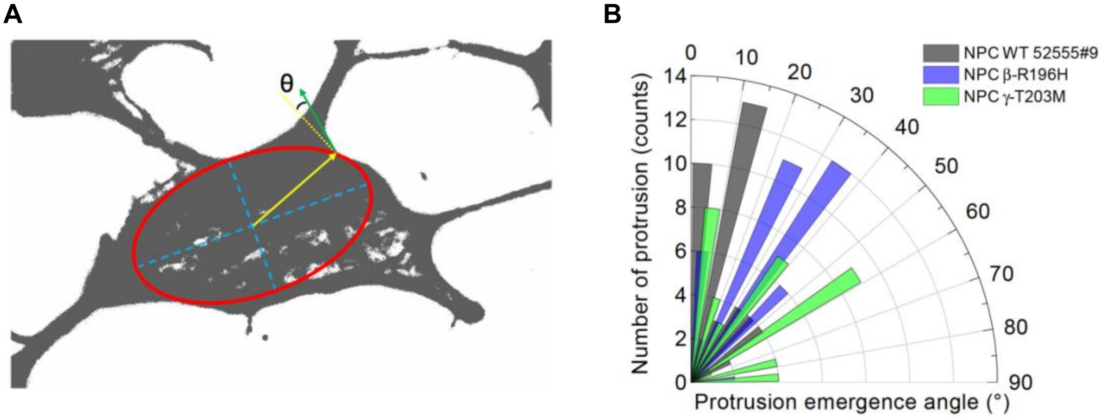

**Fig. S7. Quantification of the protrusion emergence angle relative to the soma boundary in NCPs. A.** The contour of the soma was approximated by fitting an ellipse to the cell body (red line). The center of the ellipse (blue cross) was used to define a radial vector (yellow arrow) connecting the soma center to the base of the protrusion. The main axis of the protrusion was traced to determine its orientation (green). The emergence angle (θ) was calculated relative to the local soma tangent (yellow dotted line), defined as perpendicular to the radial vector at the protrusion base (yellow continuous line). Therefore, smaller values indicate protrusions emerging with an orientation closer to the local tangent, whereas larger values indicate protrusions deviating more strongly from tangential alignment. All geometrical measurements were performed using Fiji/ImageJ. **B.** Polar histograms showing the distribution of θ. WT cells (black) display a strong bias toward small θ (0-20°), whereas β-R196H (blue) and γ-T203M (green) mutant cells show progressively broader distributions shifted toward larger angles. The mean emergence angle increases from 23 ± 19°in WT NPCs to 30 ± 18° and 40 ± 27° in β-R196H NPCs and γ-T203M NPCs, respectively. Notably, the difference between WT and β-R196H NPCs is not significant ($p = 0.072$) while the difference between WT and γ-T203M NPCs is significant ($P < 0.01$).

**Figure S8.**

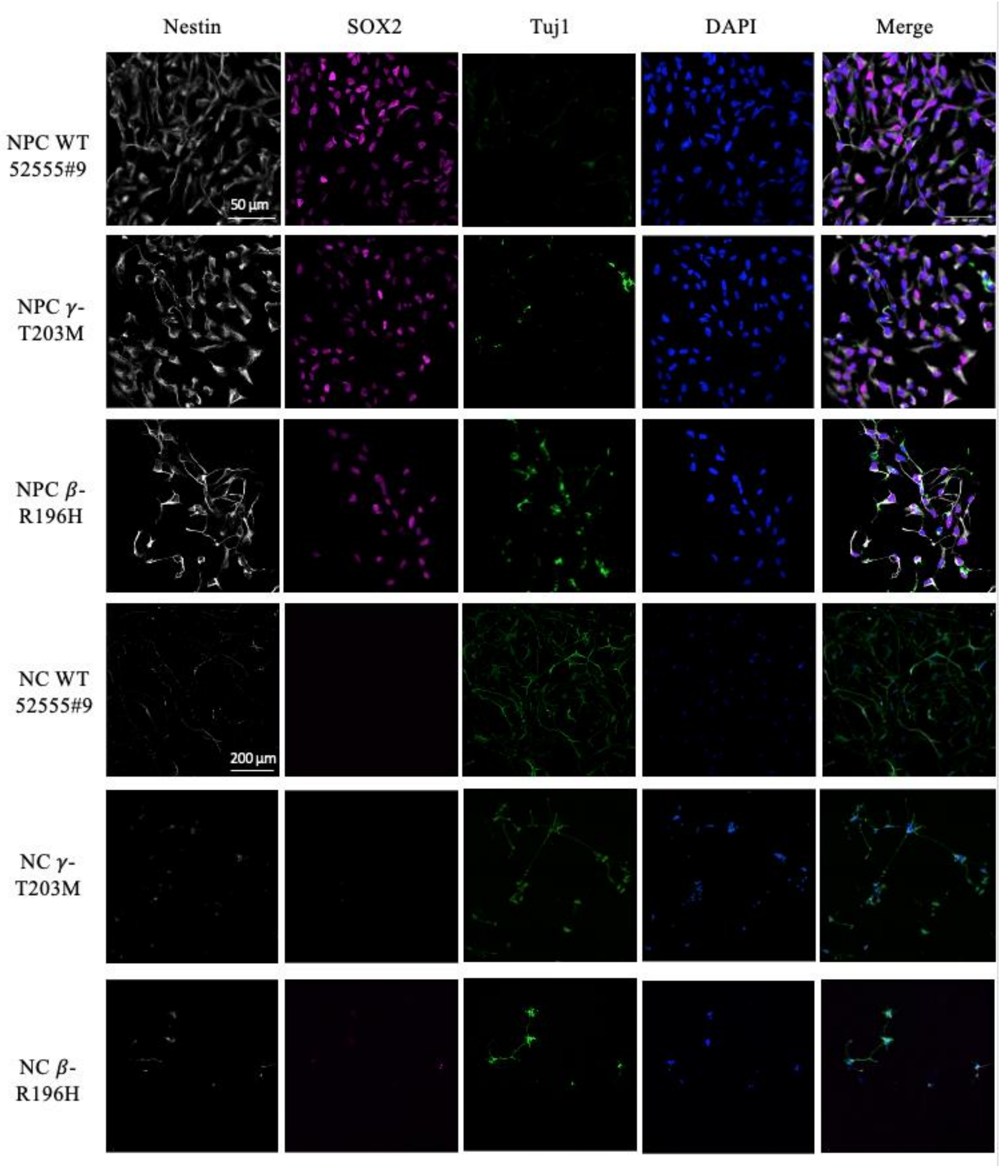

**Fig. S8**. **Comparative immunofluorescence analysis of the two maturation stages.** The comparison includes WT (52555#9), γ-T203M and β-R196H cell lines. Staining reveals expression of Nestin (grey, NPC marker), SOX2 (magenta, NPC marker), Tuj1 (green, specific for the neuron-specific βIII-tubulin isoform used as a neuronal marker), and DAPI (blue) for nuclear staining. The merged images show co-localization of the markers. Scale bars: 50 µm (NPC) and 200 µm (NC).

**Table S1. Comparison of soma size and number of cells overcoming the fixation between WT lines (52555#9 and SCTi003-A).**

| Experimental model | | Area (µm$^2$) | N | *n* | p | $N_F$ | N | *n* | p |
|---|---|---|---|---|---|---|---|---|---|
| NPC | 52555#9 | 168 ± 40 | 3 | 288 | | 91± 15 | 2 | 13 | |
| NPC | SCTi003-A | 159 ± 20 | 3 | 240 | 0.84 | 87 ± 21 | 2 | 9 | 0.87 |
| NC | 52555#9 | 620 ± 80 | 5 | 220 | | 100 ± 24 | 2 | 11 | |
| NC | SCTi003-A | 678 ± 110 | 5 | 205 | 0.67 | 96 ± 22 | 2 | 10 | 0.9 |

Morphometric analysis of soma size expressed as cell area (µm²) and of the number of cells retained on the coverslip and remaining detectable after the fixation protocol ($N_F$), expressed as mean ± SD. *N*, number of analyzed coverslips, *n*, number of cells. P values refer to comparisons between 52555#9 and SCTi003-A within the same developmental stage.

**Table S2. relevant parameters of the bi-exponential fit to force relaxation.**

| Experimental model | | *n* | $\tau_s$ (s) | $\tau_l$ (s) | $F_2$ (pN) |
|---|---|---|---|---|---|
| NPC | WT 52555#9 | 48 | 0.23 ± 0.16 | 4.6 ± 2.5 | 42 ± 4 |
| NPC | WT SCTi003-A | 21 | 0.24 ± 0.19 | 5.5 ± 3.1 | 38 ± 5 |
| NPC | γ-T203M | 11 | 0.25 ± 0.11 | 4.9 ± 1.8 | 18 ± 6 $^{\$, \#}$ |
| NPC_ | β-R196H | 53 | 0.36 ± 0.21 | 5.6 ± 3.3 | 24 ± 5 $^{\$, \#}$ |
| NPC | 52555#9 + Latr | 15 | 0.22 ± 0.11 | 3.8 ± 2.1 | 5.6 ± 2.3 |
| NC | WT 52555#9 | 20 | 0.28 ± 0.14 | 5.4 ± 2.7 | 26 ± 3 * |
| NC | WT SCTi003-A | 16 | 0.26 ± 0.09 | 5.1 ± 2.2 | 21 ± 3 * |
| NC | γ-T203M | 9 | 0.24 ± 0.12 | 4.7 ± 2.1 | 10 ± 4 $^{\#}$ |
| NC | β-R196H | 27 | 0.24 ± 0.14 | 4.6 ± 2.6 | 24 ± 4 $^{\$}$ |
| NC | 52555#9 + Latr | 25 | 0.19 ± 0.09 | 3.4 ± 1.6 | 5 ± 2 |

Parameters of the bi-exponential fit of the force relaxation time course following an elongation at 4 μm $s^{-1}$ of amplitude ~ 40 μm. *n* represents the number of tested cells. Values are expressed as mean ± SD. Symbols indicate statistically significant differences ($p < 0.05$) for the following comparisons: * WT NPCs *vs* NCs (maturation effect); $^{\$}$ Latrunculin-A-treated *vs* mutant cells; $^{\#}$ WT *vs* mutant cells.

**Table S3. Comparison of mechanical parameters between WT lines (52555#9 and SCTi003-A).**

| Experimental model | | $F_{thr}$ (pN) | | | $k_0$ (pN μm$^{-1}$) | | | $\eta_{eff}$ (pN·s μm$^{-1}$) | | |
|---|---|---|---|---|---|---|---|---|---|---|
| | | | *n* | p | | *n* | p | | *n* | p |
| NPC | 52555#9 | 15.7 ± 0.9 | 98 | | 0.53 ± 0.03 | 98 | | 0.81 ± 0.13 | 19 | |
| NPC | SCTi003-A | 14.2 ± 1.3 | 31 | 0.34 | 0.51 ± 0.04 | 31 | 0.69 | 0.86 ± 0.17 | 15 | 0.82 |
| NC | 52555#9 | 6.2 ± 1.7 | 57 | | 0.56 ± 0.07 | 57 | | 0.51 ± 0.09 | 22 | |
| NC | SCTi003-A | 4.9 ± 1.1 | 27 | 0.51 | 0.57 ± 0.04 | 27 | 0.9 | 0.49 ± 0.12 | 44 | 0.89 |

Mechanical parameters from the two WT lines, including threshold force ($F_{thr}$), tether stiffness ($k_0$), and effective viscosity ($\eta_{eff}$). Data are expressed as mean ± SD; *n* indicates the number of analyzed cells. P values indicate statistical significance between 52555#9 and SCTi003-A at the same developmental stage.

**Table S4. Tether radius for different experimental models**.

| Experimental model | | $n$ | $\sigma_a$ (nm) | $\sigma_o$ (nm) | $\sigma_t$ (nm) | ≈ $r_t$ (nm) (defined as $2\sigma_t$) |
|---|---|---|---|---|---|---|
| NPC | WT | 14 | 144 ± 3 | 155 ± 4 | 56 ± 13 | 112 ± 26 |
| NPC | γ-T203M | 7 | | 155 ± 4 | 56 ± 13 | 112 ± 26 |
| NPC | β-R196H | 8 | | 153 ± 5 | 51 ± 17 | 102 ± 35 |
| NPC | 52555#9 + Latr | 8 | | 204 ± 8 | 146 ± 13 | 292 ± 26 $^{\$}$ |
| NC | WT | 12 | | 151 ± 3 | 46 ± 15 | 92 ± 30 |
| NC | γ-T203M | 7 | | 151 ± 4 | 46 ± 16 | 92 ± 32 |
| NC | β-R196H | 7 | | 153 ± 3 | 51± 12 | 102 ± 24 |
| NC | 52555#9 + Latr | 6 | | 208 ± 7 | 151 ± 10 | 302 ± 20 $^{\$}$ |

The deconvoluted width of the tether image ($\sigma_t$) is derived from the observed Gaussian profiles ($\sigma_o$) using the system resolution $\sigma_a$ and the radius of the tether $r_t$ is taken as twice $\sigma_t$, as detailed in Materials and Methods. Values are mean ± SD. $n$ indicates the number of tethers analyzed. $^{\$}$ indicates a significant difference ($p < 0.05$) between Latrunculin-A-treated and mutant cells.

**Table S5. α-SMA expression across cellular models**

| Experimental model | | N | $n$ | $n^+$ | % |
|---|---|---|---|---|---|
| NPC | WT | 5 | 389 | 0 | 0 % |
| NPC | $\gamma$-T203M | 5 | 324 | 84 | 26 % |
| NPC | $\beta$-R196H | 5 | 262 | 0 | 0 % |
| NC | WT | 5 | 304 | 0 | 0 % |
| NC | $\gamma$-T203M | 5 | 603 | 92 | 15 % |
| NC | $\beta$-R196H | 5 | 167 | 0 | 0 % |

The table reports the number and the percentage of cells expressing α-smooth muscle actin (α-SMA) for all experimental models. Data are obtained from five images (N) per condition, by counting the total number of cells ($n$) and cells expressing α-SMA ($n^+$) in the set of images. Fisher's exact test revealed significant differences in α-SMA expression for NPC γ-T203M vs NPC WT, NPC γ-T203M vs NPC β-R196H, NC γ-T203M vs NC WT, and NC γ-T203M vs NC β-R196H ($p < 0.00001$). No significant differences were observed in the remaining comparisons ($p > 0.05$).

**Table S6. β-actin and γ-actin colocalization in NPCs and NCs.**

| | Cell stage | Cell line | PCC ($r$) | M1 | M2 | Number of measurements |
|---|---|---|---|---|---|---|
| Whole cell | NPC | WT | 0.88 ± 0.04 | 0.84 ± 0.02 | 0.82 ± 0.01 | 2 images |
| | NPC | γ-T203M | 0.94 ± 0.01 | 0.94 ± 0.03 | 0.83 ± 0.06 | 2 images |
| | NPC | β-R196H | 0.94 ± 0.03 | 0.85 ± 0.09 | 0.92 ± 0.02 | 2 images |
| | NC | WT | 0.97 ± 0.01 | 0.96 ± 0.01 | 0.93 ± 0.01 | 3 images |
| | NC | γ-T203M | 0.95 ± 0.01 | 0.94 ± 0.05 | 0.87 ± 0.08 | 2 images |
| | NC | β-R196H | 0.92 ± 0.03 | 0.81 ± 0.09 | 0.95 ± 0.06 | 3 images |
| Soma ROI | NPC | WT | 0.89 ± 0.04 | 0.88 ± 0.04 | 0.85 ± 0.02 | 42 ROIs |
| | NPC | γ-T203M | 0.94 ± 0.01 | 0.93 ± 0.02 | 0.85 ± 0.03 | 47 ROIs |
| | NPC | β-R196H | 0.94 ± 0.02 | 0.87 ± 0.09 | 0.92 ± 0.01 | 34 ROIs |
| Protrusion ROI | NPC | WT | 0.88 ± 0.04 | 0.89 ± 0.01 | 0.72 ± 0.04 | 83 ROIs |
| | NPC | γ-T203M | 0.94 ± 0.01 | 0.87 ± 0.06 | 0.88 ± 0.02 | 62 ROIs |
| | NPC | β-R196H | 0.94 ± 0.02 | 0.83 ± 0.09 | 0.99 ± 0.01 | 59 ROIs |

Pearson correlation coefficient (PCC, $r$) and Manders coefficients (M1 and M2) were calculated from fluorescence images of WT, γ-T203M, and β-R196H cells (see Materials and Methods). Analyses were performed on whole-cell images for NPCs and NCs and on regions of interest (ROIs) corresponding to the soma and to intercellular protrusions in NPCs. ROI analyses were performed on the same images used for whole-cell measurements. Values represent mean ± SD.

**Table S7. Primary antibodies used for immunofluorescence.**

| Target | Company | Catalog number | Source | Isotype | Species reactivity | Cellular localization | Dilution |
|---|---|---|---|---|---|---|---|
| β-actin | Bio-Rad | MCA5775GA | Mouse monoclonal | IgG1 | Human | Cytoskeleton | 1:100 |
| γ-actin | Bio-Rad | MCA5776GA | Mouse monoclonal | IgG2b | Human | Cytoskeleton | 1:200 |
| α-smooth muscle actin | abcam | ab7817 | Mouse monoclonal | IgG2a | Human | Cytoskeleton | 1:500 |
| TUBB | abcam | ab179513 | Rabbit monoclonal | IgG | Human | Cytoskeleton | 1:500 |
| Sox2 | R+D Systems | AF2018 | Goat polyclonal | IgG | Human | Nucleus | 1:300 |
| Nestin | Sigma-Aldrich | N5413-100UG | Rabbit monoclonal | IgG | Human | Intermediate filaments | 1:300 |
| Tuj1 | BioLegend | 801202 | Mouse monoclonal | IgG2b | Human | Microtubules | 1:300 |

List of primary antibodies used for immunofluorescence, including target, company, catalog number, host species, isotype, species reactivity, cellular localization, and working dilution.

**Table S8. Secondary antibodies used for immunofluorescence.**

| Target of the primary antibody | Company | Catalog number | Type | Source | Isotype | Conjugate | Dilution |
|---|---|---|---|---|---|---|---|
| β-actin | Jackson IR | 115-545-205 | Anti-mouse | Donkey | IgG1 | Alexa Fluor® 488 | 1:200 |
| γ-actin | Jackson IR | 115-175-207 | Anti-mouse | Donkey | IgG2b | Cyanine Cy™5 | 1:50 |
| α-smooth muscle actin; Tuj1 | Thermo Fisher | A-21202 | Anti-mouse | Donkey | IgG | Alexa Fluor® 488 | 1:500 |
| TUBB; Nestin | Thermo Fisher | A-31572 | Anti-rabbit | Donkey | IgG | Alexa Fluor® 555 | 1:500 |
| Sox2 | Thermo Fisher | A-21447 | Anti-goat | Donkey | IgG | Alexa Fluor® 647 | 1:500 |

List of secondary antibodies used for immunofluorescence, including target, company, catalog number, host species, isotype, species reactivity, cellular localization, and working dilution.

**Table S9. Primary antibodies used for Western blotting**

| Target | Company | Catalog number | Source | Isotype | Species reactivity | Dilution |
|---|---|---|---|---|---|---|
| β-actin | Bio-Rad | MCA5775GA | Mouse monoclonal | IgG1 | Human | 1:750 |
| γ-actin | Bio-Rad | MCA5776GA | Mouse monoclonal | IgG2b | Human | 1:7500 |
| α-smooth muscle actin | abcam | ab7817 | Mouse monoclonal | IgG2a | Human | 1:300 |
| GAPDH | Thermo Fisher, Invitrogen | MA5-15738 | Mouse monoclonal | IgG | Human | 1:1000 |
| Pan-Actin | Cell Signaling | 8456 | Rabbit monoclonal | IgG | Human | 1:1000 |

Primary antibodies used for Western blotting and their specifications, including target protein, company, catalog number, host, and isotype, species reactivity, and working dilution.

**Table S10. Secondary antibodies used for Western blotting**

| Target of the primary antibody | Company | Catalog number | Type | Source | Isotype | Conjugate | Dilution for WB |
|---|---|---|---|---|---|---|---|
| β-actin, γ-actin, α-smooth muscle actin, GAPDH | Merck | 12-349 | Anti-mouse | Goat | IgG | HRP conjugate | 1:10000 |
| Pan-Actin | Sigma-Aldrich | 12-348 | Anti-Rabbit | Goat | IgG | HRP conjugate | 1:2000 |

Secondary antibodies used for Western blotting, including target of the primary antibody, company, catalog number, antibody type, host species, isotype, conjugate, and working dilution.
